# Oblique Line Scan Illumination Enables Expansive, Accurate and Sensitive Single Protein Measurements in Solution and in Living Cells

**DOI:** 10.1101/2023.12.21.571765

**Authors:** Amine Driouchi, Mason Bretan, Brynmor J. Davis, Alec Heckert, Markus Seeger, Maité Bradley Silva, William SR. Forrest, Jessica Hsiung, Jiongyi Tan, Hongli Yang, David T. McSwiggen, Linda Song, Askhay Sule, Behnam Abaie, Brianna Conroy, Liam A. Elliott, Eric Gonzalez, Fedor Ilkov, Joshua Isaacs, George Labaria, Michelle Lagana, Delaine D. Larsen, Brian Margolin, Mai K. Nguyen, Jeremy Rine, Yangzhong Tang, Martin Vana, Andrew Wilkey, Zhengjian Zhang, Stephen Basham, Jaclyn J. Ho, Stephanie Johnson, Aaron A. Klammer, Kevin Lin, Xavier Darzacq, Eric Betzig, Russell T. Berman, Daniel J. Anderson

**Affiliations:** Eikon Therapeutics; EikonTherapeutics; UC Berkeley; Department of Molecular & Cell Biology, University of California, Berkeley, Berkeley, CA, USA

## Abstract

Single-molecule localization microscopy (SMLM) techniques, such as single-molecule tracking (SMT), enable *in situ* measurements in cells from which data-rich metrics can be extracted. SMT has been successfully applied to a variety of biological questions and model systems, aiming to unravel the spatiotemporal regulation of molecular mechanisms that govern protein function, downstream pathway effects, and cellular function. While powerful, SMLM often suffers from low throughput and illumination inhomogeneity, along with microscope and user-induced technical biases. Due to technical limitations in scaling SMLM techniques, a tradeoff between spatiotemporal resolution and throughput has been made historically, restricting broad application of these technologies. Here we address these limitations using Oblique Line Scan (OLS), a robust single-objective light-sheet based illumination and detection modality that achieves nanoscale spatial resolution and sub-millisecond temporal resolution across a 250 × 190 μm field of view. We demonstrate OLS-enabled SMT on Halo-Tagged proteins in living cells capturing protein motion up to 14 μm^2^ /s. By exploiting the adaptability of the acquisition frame rate and the improved rejection of out of focus light, we extend the utility of OLS beyond cellular compartments with in-solution SMT (isSMT) for single-molecule measurement of ligand-protein interactions and disruption of protein-protein interactions (PPI). We illustrate the versatility of OLS by showcasing two-color SMT, STORM, and single molecule fluorescence recovery after photobleaching (FRAP). OLS expands the range of SMLM applications and paves the way for robust, high-throughput single-molecule investigations of protein dynamics required for drug screening and systems biology studies, both in cells and in solution.

## Introduction

A protein’s function within a living cell is controlled by a myriad of factors defined by its specific physiological context and dynamic local environment. While a given protein’s role is tightly linked to its structure, expression levels, self-association states and interaction partners, a complete understanding of the underlying biological complexity requires measurements that provide detailed spatiotemporal context. Several technological advancements have given rise to super-resolution microscopy techniques capable of circumventing the diffraction limit of light, hence enabling the localization of individual proteins in cells. SMLM comprises a subset of super-resolution microscopy methods that rely on detecting and localizing sparse fluorescently labeled proteins (Chi 2009; Liu et al., 2015). Such SMLM techniques include stochastic optical reconstruction microscopy (STORM) (Rust et al., 2006) and photoactivation localization microscopy (PALM) (Betzig et al., 2006; Shroff et al., 2008). Both techniques rely on the stochastic photoswitching or photoactivation of organic fluorophores or fluorescent proteins, respectively, to achieve individual point spread function (PSF) localization.

Capturing individual or ensemble protein dynamics in living cells at single-molecule resolution also relies on sparse fluorescent labeling of proteins to facilitate sub-diffraction-limit PSF localization. In addition, single molecule tracking (SMT) requires adequate temporal sampling to enable computational frame-to-frame linking and trajectory generation from which a diffusion coefficient can be calculated (Levi and Gratton 2007). Due to the relatively low quantum yields and extinction coefficients of previously used fluorophores, early implementations of SMT predominantly focused on motor protein motion along microtubules (Yildiz et al., 2004) or 2D protein dynamics at the plasma membrane (Ewers et al., 2005; Jin et al., 2007; Cui et al., 2018), which are intrinsically slow processes that can be sampled with long camera exposure times. The advent of highly sensitive sCMOS and EMCCD cameras, the use of stroboscopic illumination to reduce motion-induced blurring, and the development of superior labeling strategies in conjunction with bright, live-cell compatible fluorophores have enabled SMT measurement of intracellular, fast and transient processes such as transcription factor association and dissociation at promoter region sites using short exposure times (Elf et al., 2007; Los et al., 2008; Huang et al., 2013; Izeddin et al., 2014; Grimm et al., 2015).

However, several of these advanced microscopy techniques suffer from low throughput, uneven illumination, and instrument- and user-induced biases such as manual field of view (FOV) selection. Traditional through-the-objective illumination schemes designed to minimize out-of-focus background, such as Highly Inclined and Laminated Optical Sheet (HILO) are easily implemented, requiring only a high numerical aperture (NA) objective, and are compatible with commonly used sample holders and imaging chambers (Tokunaga et al., 2008). Unfortunately, HILO methods still suffer from uneven illumination, forcing the capture of a smaller region of the camera chip to reduce detection and reconstruction artifacts when used in SMLM or SMT. A variety of illumination schemes have been employed to remediate this uneven illumination, such as the azimuthal beam scanning that is used in several applications of Total Internal Reflectance Fluorescence Microscopy (i.e., spinning TIRF) (Zong et al., 2014; Deschamps et al., 2016). While these methods improve signal-to-noise ratio (SNR) at the glass-sample interface, they do not provide sizable improvements in rejecting out-of-focus light hence negatively impacting overall SNR. Light-sheet-based illumination strategies have been developed that provide superior optical sectioning capabilities, larger captured FOV and homogeneous sample illumination. Lattice light-sheet microscopy, a modified version of light-sheet microscopy utilizing a structured light sheet, has been demonstrated to provide such improvement. While Lattice light-sheet microscopy and other multi-objective light-sheet approaches have demonstrated high performance in capturing single-molecule measurement in a variety of live and fixed samples, the necessity for 2 objectives, where one objective is used to illuminate the sample and the other to collect light, impedes implementations and service at scale. While powerful, this geometry also restricts the type and size of samples that can be mounted between the objectives for imaging. Other light-sheet based microscopy modalities have since been developed aiming to avoid specialized optics and customized sample holders. Selective plane illumination microscopy utilizes micromirrors to generate a light sheet that can be captured through a single objective and has been applied to capture SMLM datasets (Galland et al., 2015). Single-molecule oblique-plane microscopy has demonstrated SMLM capabilities within thick samples via the use of a water-immersion objective and polarization optics (Kim et al., 2019). More recently, single-objective lens inclined light sheet and high-contrast single-molecule imaging by highly inclined swept tile illumination have combined oblique illumination, PSF engineering, and light-sheet translation to achieve FOV acquisition of 25 × 45 μm and 130 × 130 μm, respectively (Tang and Han 2018; Hung et al., 2022).

Here, we introduce OLS, a robust light-sheet-based illumination strategy that enables the capture of a FOV six-fold larger (250 × 190 μm) than what is typically captured in HILO, within a 384 well-plate, with nanoscale spatial resolution and sub-millisecond temporal resolution. The relative simplicity of the optical configuration used in OLS renders this approach readily implemented on inverted microscopes equipped with either water- or oil-immersion high numerical aperture (NA) objectives and a sCMOS camera with light-sheet mode capability. We have applied this modality to *in situ* live-cell SMT on protein targets of interest. The unique setup and the fully automated acquisition scheme implemented on OLS microscopes simplifies usability and maximizes throughput and versatility, importantly, while overcoming some of the key limitations of previously described work focusing on light-sheet based SMLM. Using OLS-enabled fast SMT, optimized tracking algorithms, along with machine learning-based segmentation models, we demonstrate the system’s suitability for interrogating and capturing the baseline biological heterogeneity and cell cycle dependency of the proliferating cell nuclear antigen (PCNA). Our adjustable and high frame rate, along with our homogenous illumination, ultimately enable in isSMT to study protein-protein and protein-ligand interactions, demonstrated by capturing single protein dynamics of TCF4/Beta-Catenin. Finally, we present the breadth of SMLM-based methods and advanced microscopy techniques that can be successfully applied using OLS microscopy. We highlight the suitability of our system to conduct two-color SMT and FRAP. Overall, OLS enables large scale, robust single-molecule imaging, paving the way to start exploiting novel mechanistic insights derived from single-molecule measurements towards a deeper understanding of the underlying biology important to human disease and drug discovery applications.

## Results

### OLS provides near full field homogeneous illumination enabling SMT

OLS was developed with the purpose of enlarging the effective imaging area while homogenizing SNR across the camera chip to yield high quality SMLM and SMT raw image files with no compromise to the achieved spatiotemporal resolution. Briefly, a thin optical light-sheet is shaped and focused into the back focal plane of the microscope’s objective and scanned using a galvanometric mirror. This optical configuration results in a scannable oblique light sheet that can cover the full FOV of the water-immersion high NA objective (Figure 1A and S1). We characterized the performance of this imaging system, leveraging a U2OS cell line with an N-terminally Halo-Tagged KEAP1 construct (Halo-KEAP1). KEAP1 provides a simple model system with which, SMT performance can be characterized. Our initial imaging of Halo-KEAP1 sparsely labeled with the rhodamine dye Janelia Fluorophore 549 (JF_549_) resulted in clear single molecule resolution data suitable for the analysis of spot detection, localization and tracking of individual spots (Figure 1B and Methods). We benchmarked the performance of the OLS system against a HILO implementation on the same microscope (see Methods). 1.5 seconds of SMT data were collected in both HILO and OLS and the resulting trajectories of Halo-KEAP1 were plotted (Figures 1D). The average number of trajectories collected across the FOV increased from 25,765 ± 4,838 with HILO to 167,479 ± 46,324 with OLS, matching the calculated 6-fold imaging field increase (Figure 1E). SMT data was collected for 1,224 FOVs across a 384 well plate and the average SNR for all spots localized within each pixel of the FOV was calculated and a spatial SNR map was rendered (Figure 1F). The standard deviation and average SNR per FOV were then summarized across 308 wells for OLS and HILO, and clearly demonstrate improved consistency and performance in SNR of OLS when comparing the two illumination modalities (Figure 1G).

**Figure 1:**
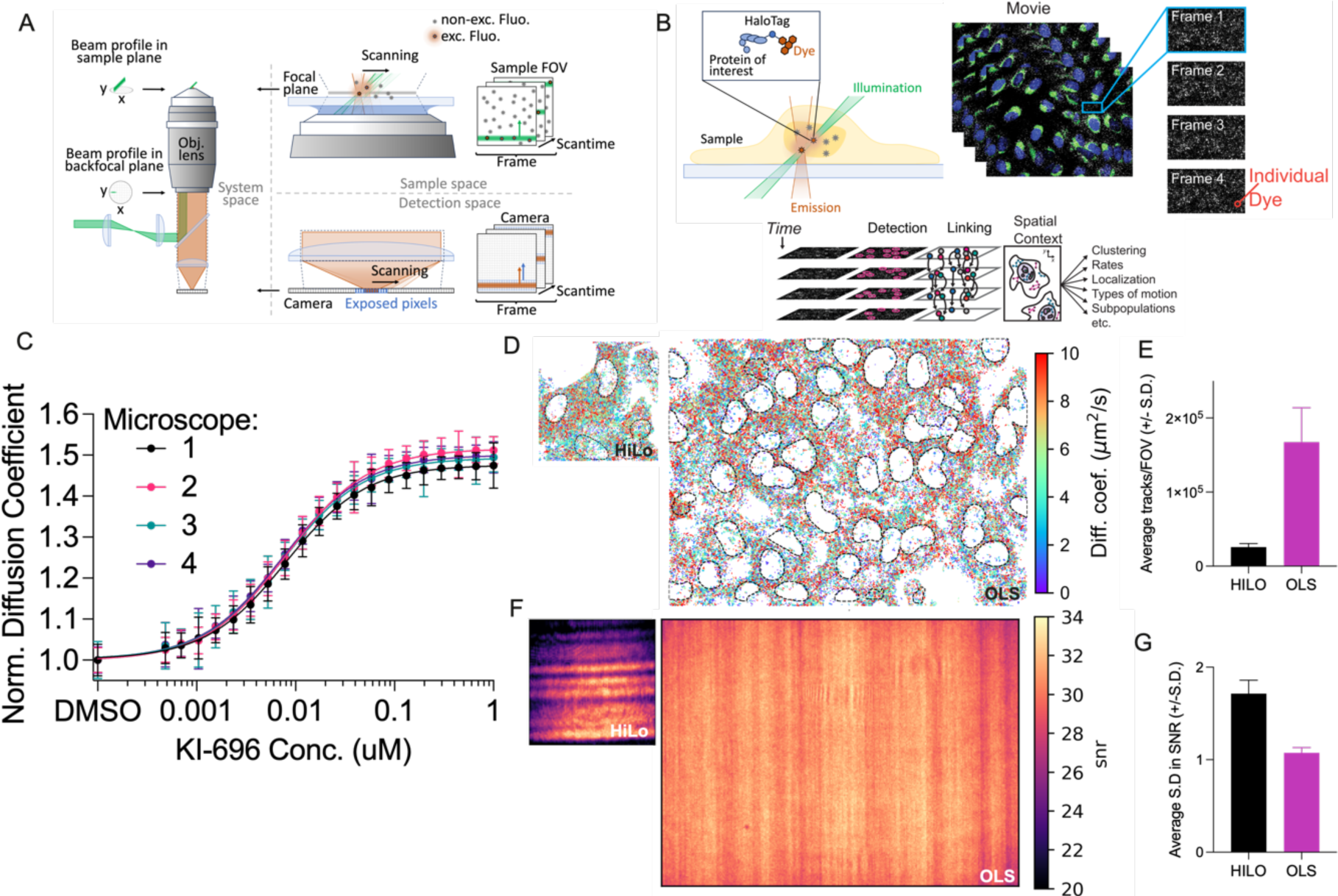
OLS provides near full field homogeneous illumination enabling SMT. **(A)** Schematic representation of the OLS implementation. A collimated beam is shaped into an optical light-sheet, which is sent to a water-immersion objective with the emission being projected onto a high-speed sCMOS camera. **(B)** The SMT workflow. SMT setup relies on the Halo-tagging of protein targets of interest. JF_549_ or JF_646_ organic fluorophores are used to provide individual emitters with appropriate signal to conduct frame to frame linking and track generation. From these coordinates and trajectories, a variety of metrics including protein diffusion and spatial localization can be extracted. **(C)** OLS generated dose-response curves of the Keap1 inhibitor KI-696 on U2OS cells expressing halo-tagged Keap1. Data was acquired across 4 microscopes and imaged on our high throughput SMT platform. 72 FOVs from 12 wells were captured for each concentration on randomized plates, error bars denote standard deviation. **(D)** Representative sampling areas imaged with HILO and OLS illumination from Halo-Keap1 expressing U2OS cells. Trajectories are plots across a 1.5 s acquisition and color coded based on the measured diffusion coefficient with nuclear mask outlines overlaid with a black dotted line. **(E)** Quantification of number of trajectories captured per FOV using HILO and OLS. **(F)** Representative average spatial SNR maps per pixel calculated across 1,232 FOVs for a plate imaged with either HILO or OLS. **(G)** Average FOV-level standard deviation in SNR sampled over 308 wells for HILO or OLS illumination.

Variations in the performances of different microscopes can drastically limit interpretations of consistency, robustness and applicability of implementation. To evaluate the reproducibility in illumination quality of the OLS optical systems, we conducted side-by-side SMT measurements on four distinct OLS-equipped microscopes using our previously described automated system (McSwiggen *et al*., 2023). For these experiments, we continued to use the Halo-KEAP1 U2OS cell line along with KI-696, a small molecule inhibitor known to disrupt the interaction of KEAP1 with its binding partner NRF2. Inhibition of KEAP1/NRF2 binding results in an increase in the monomeric fast-diffusing fraction of Halo-KEAP1 in this cell line, observable by SMT. Halo-KEAP1 cells were treated in 384-well plates with a randomized 20-point dose-response of KI-696 testing 12 wells per concentration and six FOVs per well. The average dose-response profiles per microscope were consistent, with a median increase in diffusion of 47-51%, and resulting median EC_50_ values ranging between 7.37 and 8.58 nM across the 4 independent microscope setups (Figure 1C and S2A). We compared the average FOV-level SNR per microscope and observed that all 4 microscopes provided a median SNR ranging between 28.08 and 28.89 (Figure S2B). Importantly, we did not observe changes across subsequent FOVs captured within a single well, suggesting minimal disruption within a well upon imaging (Figure S2C). Additionally, we directly characterized the effect of the large OLS FOV size on SMT sampling by comparing cropped regions of the same FOV to the large OLS sized-FOV. We observed a significant increase in the variance as we shrink the number of captured cells to an area spanning 83 × 83 μm (Figure S2D), highlighting the importance of a sufficiently large FOV to adequately capture the intrinsic cell to cell variability of a sample.

Given the demonstrated improvement in OLS SNR, we next wanted to explore other improvements in system performance. For a typical HILO implementation, samples are illuminated for 2 ms by pulsing the excitation laser for a subset of the camera exposure time. For OLS, given the scanning rate of the light-sheet, we calculated that each fluorophore is only exposed to light for 400 μs. Given this shorter illumination time, we expected more consistent point spread functions (PSFs) across different diffusion rates. We tested this hypothesis by analyzing mean spot width of Halo-KEAP1 with and without KI-696 (Figure S3A). With HILO illumination, there is a 4.4% increase in the mean 2σ radius of single PSFs which is decreased to 1.4% for OLS (Figure S3B and D). While a 400 μs strobe time for HILO would have provided a direct comparison in motion-induced blurring performance with OLS, we find that with this illumination time, HILO does not enable single molecule detection, as the vast majority of PSFs do not pass the noise threshold (Figure S3C), further highlighting the improved sensitvity of OLS illumination.

The impact of optical sectioning performance on an imaging system can be significant. A major advantage provided by OLS is that, during the scan of the inclined light sheet, out of focus illuminated emitters reside outside of the recorded strip of pixels on the camera. We characterized this illumination-based optical sectioning method by preparing increasing concentrations of His-HaloTag in solution to titrate protein labeling density and the downstream effect on SNR and PSF detection. This experiment captures the expected improvement in the sectioning ability provided by OLS, where we observed a faster decrease in the number of detected localizations in HILO, which is correlated to a dye concentration-dependent decrease in SNR (Figure S3E and F). These results highlight that under OLS illumination, we can better detect single PSFs irrespective of local PSF overlaps that may arise from increasing dye or protein concentration. Taken together with the reduction in motion blur, OLS provides an improvement in the ability to track single particles at high density with high resolving performance. Combined, these results highlight the robustness and reproducibility of SMT measurements conducted using OLS illumination within a large FOV, and demonstrate the superior performance of OLS compared to HILO when characterizing the motion of fast-moving proteins at high labeling density.

### OLS allows for rapid SMT data capture that enables tracking of fast-moving proteins

Having established the robustness of OLS illumination and given the line-scanning modality of OLS (Figure S1), we investigated the impact of higher frame rate acquisition on assay window improvement and key imaging metrics. We hypothesized that, given the range and specificity of protein motion in live cells, there exists a set of optimal acquisition parameters for a given protein of interest. Frame rate correlates with other experimental factors such as localization error and tracking error to determine the information recoverable from SMT (Figure S4A and B). To understand these effects, we performed optical-dynamical simulations with complex mixtures of Brownian motions (Figure S4C), then ran tracking on these simulated movies. Both mean track length and tracking fidelity improved with increasing frame rate, highlighting that the sampled FOV size is the only apparent compromise (Figure S4D). We then estimated the underlying dynamical model for each simulation using state array analysis, a variational Bayesian method for recovering mixture models from observed trajectories (Heckert et al., 2022). Increasing frame rate improved recovery of faster states but eventually degraded the recovery of slower ones (Figure S4E). We also noted that the lower and upper bounds on the mean squared displacement (MSD) estimator of the diffusion coefficient, determined on one hand by localization error and on the other by the search radius used in tracking, roughly approximated this dynamic range (Figure S4E, green dotted lines). These results suggested that tunable frame rates are a highly desirable property of an SMT imaging system.

Turning to our experimental Halo-KEAP1 system, we ran OLS-enabled SMT with frame rates ranging from 100 to 1250 Hz (Figure 2A). Similar to simulation results, both mean trajectory length and the estimated linking precision improved at higher frame rates (Figure S5A, S5C). Moreover, the OLS illuminator achieved this without degrading mean SNR (Figure S5B) or bleaching rate per frame Figure S6). Upon running state array analysis, we recovered faster motion with increasing frame rate until the estimates stabilized at ∼9 µm^2^/s for DMSO-treated Halo-KEAP1 and ∼14 µm^2^/sec for KI-696-treated Halo-KEAP1 at 400 Hz (Figure S5D, Figure 2B). Interestingly, 400 Hz may represent a point of diminishing return where the sampling frequency may be appropriate to capture the faster diffusing subset of Halo-KEAP1 under both DMSO and KI-696 treatment (Figure 2B). Further, when we simulate the measured diffusion coefficients for a KEAP1 monomeric state at varying frame rates, simulations closely matched measured results of KI-696 treated cells (Figure 2C and S4E). Together, these results demonstrate that the ability to increase frame rate utilizing the line scanning of OLS facilitates the accurate measurement of fast protein diffusion in the cellular environment. While 400 Hz appears to be an appropriate sampling speed for KEAP1, we anticipate that biochemical processes exist in live cells that will require significantly higher frame rates, making the ability to tune OLS sampling speed a desirable feature of the imaging system.

**Figure 2:**
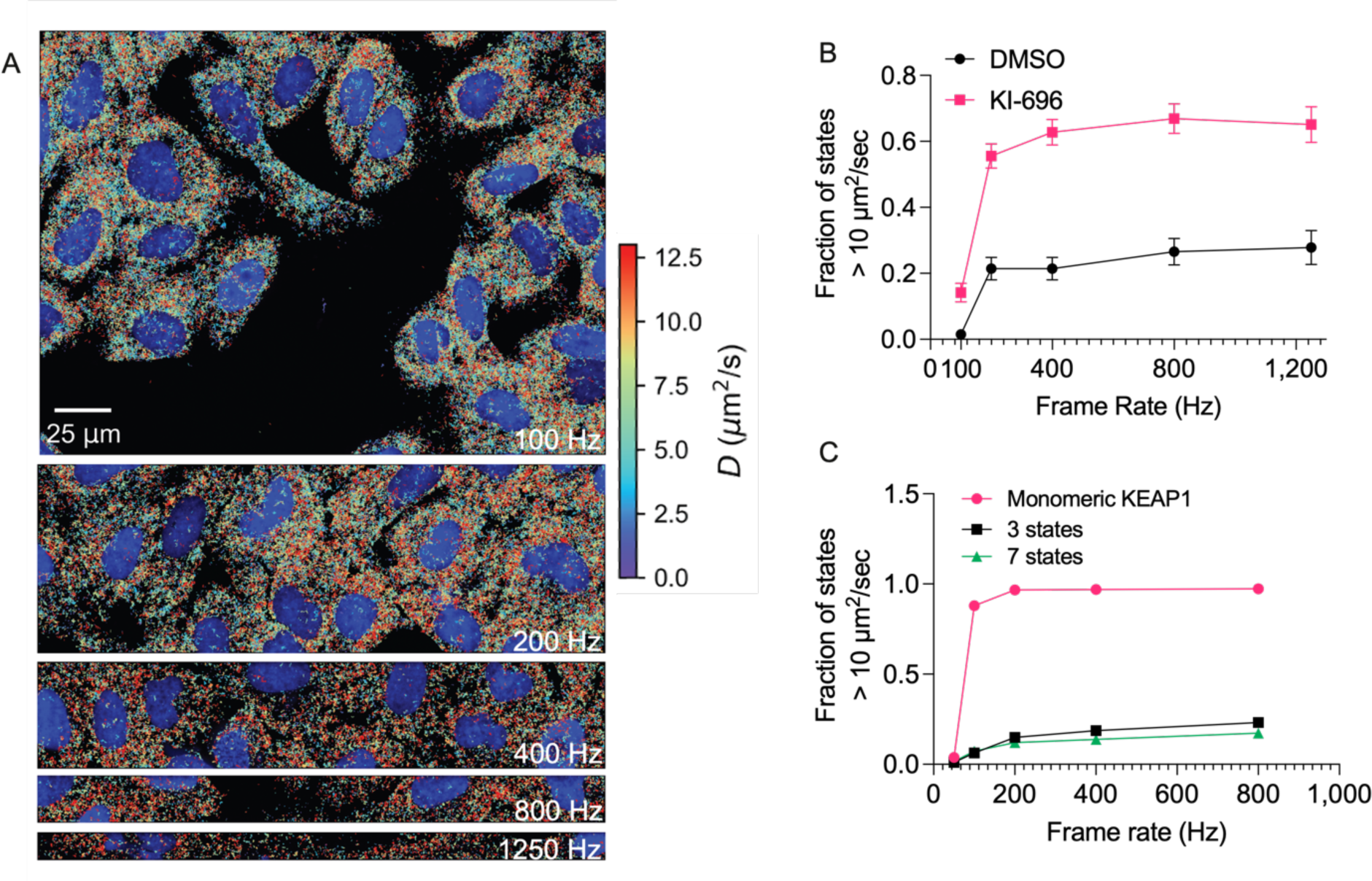
OLS enables the capture of fast protein diffusion in living cells. **(A)** Representative images of FOV size for each of five frame rates ranging from 100 - 1250 Hz. Trajectories are overlaid onto a mean projection for the Hoechst channel (blue) and colored by their maximum likelihood diffusion coefficients. **(B)** Fraction of trajectories with a diffusion coefficient > 10 μm^2^/s as function of frame rate for DMSO and KI-696-treated cells, computed from the state array posterior mean occupations. **(C)** Accuracy of state profile recovery from optical-dynamical simulations of SMT across several frame rates for 3 distinct state mixtures. Error bars denote S.D.

### OLS can be applied to capture the intercellular heterogeneity of single protein dynamics

SMT can be scaled in multiple dimensions: more trajectories per cell, more cells per FOV and more FOVs per well. Choosing optimal dimensions for scaling requires understanding which sources of variability limit the precision of downstream measurements. In assessing the consistency of SMT measurements (see Methods), we determined that intercellular heterogeneity is the highest source of variance, exceeding FOV, well, plate or microscope-level variation. We found that cell-to-cell biases were at least an order of magnitude greater than FOV-to-FOV or well-to-well biases (Figure S7). This strongly indicates that biological heterogeneity is dominant over technical variation of OLS-based SMT measurements (Figure 3A). Examples of cell heterogeneity include cell cycle state, state of cell stress, and genetic variation. Single cell measurements such as large-FOV SMT can enable more nuanced measurements to better understand such heterogeneity (Altschuler and Wu 2010). We set out to exemplify such single-cell analysis by measuring the effect of cell cycle on PCNA protein dynamics.

**Figure 3:**
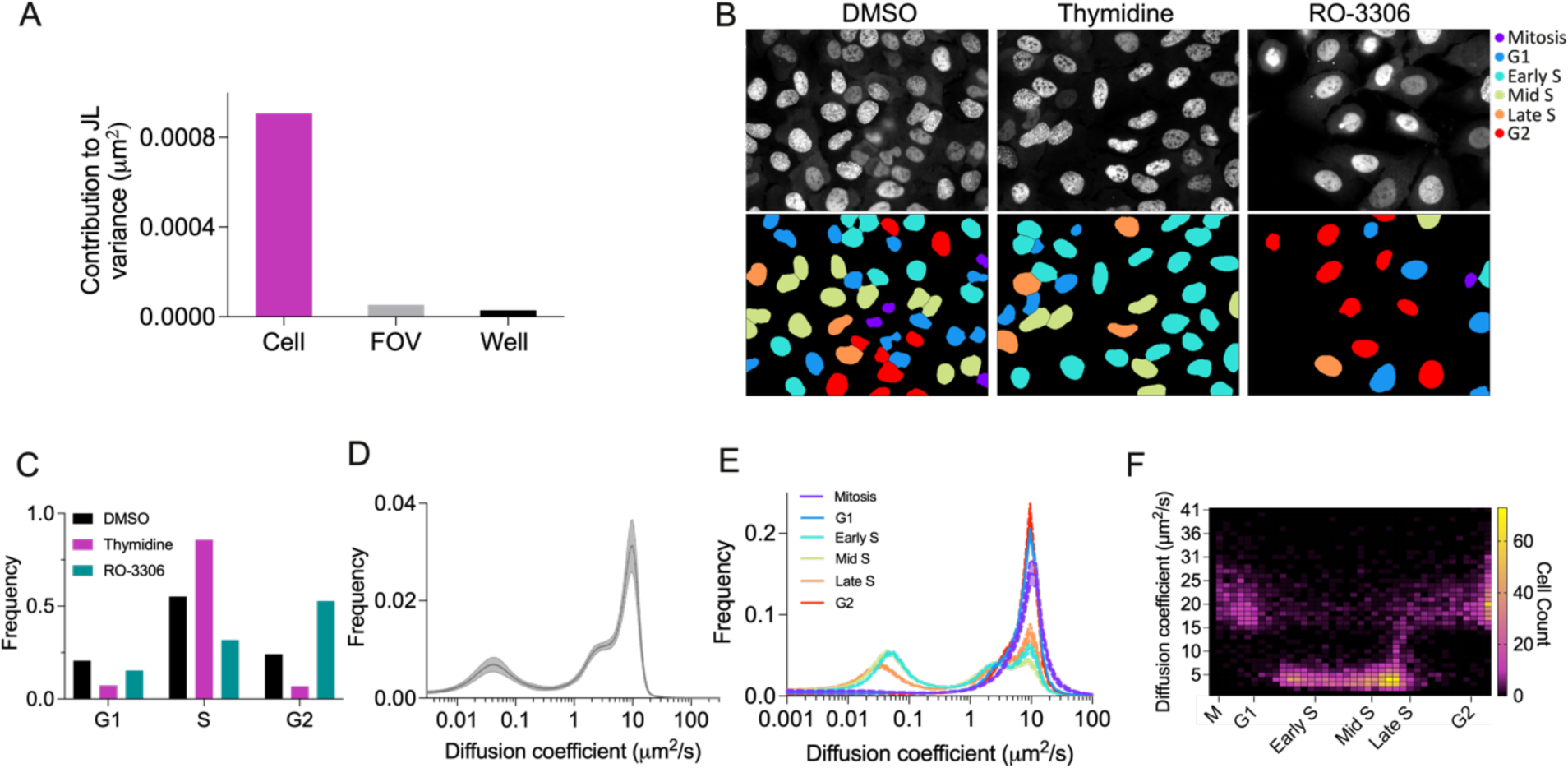
OLS can be applied to capture the inter and intracellular heterogeneity of single protein dynamics. **(A)** Analysis of sources of variance in SMT for KEAP1 measured under OLS illumination. **(B)** Representative images of Halo-PCNA labeled cells treated with 2 mM Thymidine or 10 μM RO-3306 (top). Cell cycle prediction from a machine learning (ML) model used to color cell by cycle phase (bottom). **(C)** Quantification of fraction of cells in each cell phase in response to cycle block treatments in (B). The following number of cells were analyzed for each condition: 34,067 for DMSO, 3,831 for RO-3306 and 6,044 for thymidine. **(D)** State array analysis of total population of sparsely labeled PCNA cells. **(E)** State array analysis for cells in each phase as predicted by ML model. **(F)** Heat map of 4,081 Individual cells classified using a continuous classification score plotted against PCNA diffusion coefficient.

PCNA is involved in DNA replication and localizes to replication foci during S-phase therefore exhibiting distinct and specific protein dynamics across stages of the cell cycle (Zessin et al., 2016). We introduced Halo-PCNA into U2OS isolated clones, resulting in sub-endogenous expression levels of tagged protein (Figure S8A and B) and found growth rates were not measurably affected by the expression of Halo-PCNA (Figure S8C). Proper localization of Halo-PCNA was confirmed with colocalization analysis with an RFP-labeled anti-PCNA CCR nanobody (Figure S8D).

Time-lapse microscopy was conducted on Halo-PCNA labeled with 50 nM JFX_650_, to achieve near-saturating labeling, for 12 hours at 12 frames per hour (Figure S9A). A machine learning model was then trained using both manually assigned cell stages and time-based progression resulting in both a G1, early S, mid S, late S, G2 and mitosis classifier as well as a regression prediction across the continuum of the cell cycle (Figure S9B). The predicted cell cycle classification performed well as compared to manual annotation (Figure S9C). We also observed clear progression of individual cells through the cell cycle when regression prediction was plotted across the time series (Figure S9D). To further validate our cell cycle phase assignment model, we blocked cell cycle progression at S-phases with thymidine or the G2-M transition with the CDK-1 inhibitor RO-3306 (Vassilev 2006; Chen and Deng 2018). Both treatments lead to expected enrichment in the fraction of cells assigned to the respective cell phases (Figure3B and C). Cells were then labeled with 10 pM JF_549_ and 50 nM JFX_650_ to enable simultaneous cell cycle assignment based on near-saturating labeling along with SMT measurements.

Two peaks of mobility were observed at 0.043 and 9.88 μm^2^/s, likely representing PCNA associated at sites of DNA replication and free PCNA, respectively (Figure 3D). When cells in each cell phase were analyzed separately, it became apparent that the slow-moving population of PCNA was exclusively found in cells predicted to be in S-phase and that this subset of cells also showed a marked decrease in the fast-moving population (Figure 3E). The mean diffusion coefficient of PCNA was then measured for 4,081 individual cells to characterize heterogeneity across the cell population, which is described in large part by predicted cell cycle state (Figure 3F). Taken together these data provide an example of how large-FOV SMT enabled by OLS allows for the capture and elucidation of cell-level heterogeneity of protein dynamics.

### OLS enables measurement of protein diffusion in solution

Due to the higher achievable frame rates and superior optical sectioning performance, we next wanted to test the suitability of OLS to measure the diffusive properties of purified proteins in solution. One could imagine such dynamics would be impacted by protein conformation, ligand interactions, or protein-protein interactions. As modeled by the Stokes-Einstein equation, the movement of a particle is defined by the temperature and viscosity of the medium and the effective hydrated radius of the particle (Miller 1924). The effective hydrated radius is influenced by protein size, structure, and physicochemical features of the protein surface. The diffusion of JF_549_-labeled His-Halo was measured with increasing concentrations of glycerol to modulate viscosity. A diffusion coefficient was estimated from tracking results and corrected using analytical expressions for tracking biases, resulting in a close match to the theoretical values predicted by the Stokes-Einstein equation (Figure 4A and B and see Methods).

**Figure 4:**
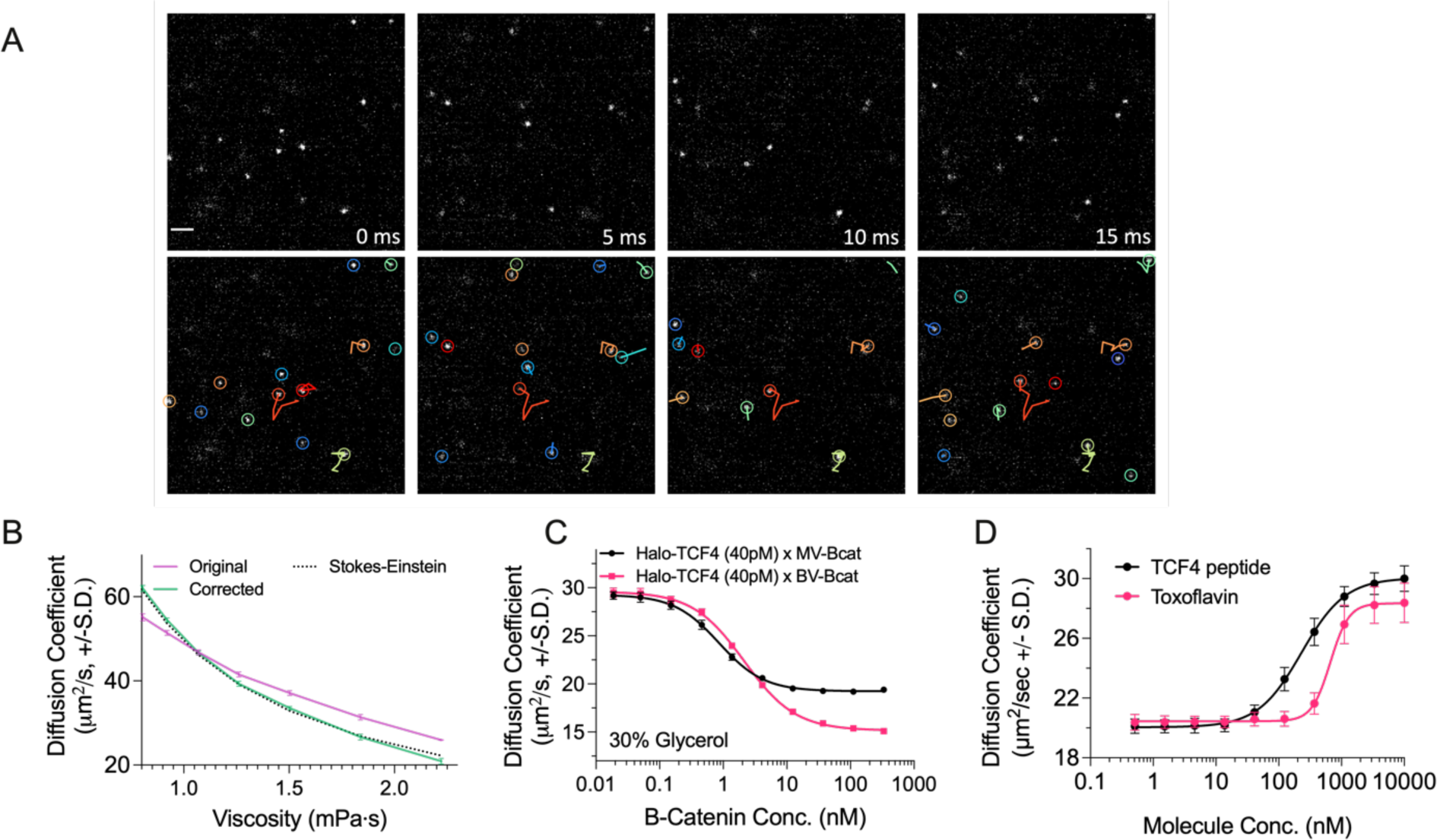
OLS enables in solution measurement of protein diffusion. **(A)** Representative sequential images of isSMT (top) with overlaid tracks (bottom) for JF_549_-labeled free HaloTag protein at 30% glycerol for a 192 × 192 pixels ROI at 200 Hz. Scale bar represents 2 μm. **(B)** Mean diffusion coefficient as a function of medium viscosity compared to theoretical Stokes-Einstein equation. **(C)** Halo-TCF4 diffusion measured with a dose response of either monomeric (monovalent/MV) vs dimeric (bivalent/BV) β-catenin protein, n= 8 wells. **(D)** Diffusion coefficient of the TCF4/β-catenin complex measured across a dose response of unlabeled TCF4 peptide and Toxoflavin, n= 32 FOVs from two distinct plates. Error bars denote S.D.

Next, we sought to measure the change in diffusion mediated by protein-protein interactions (PPI). Transcription factor 4 (TCF4) (41.5 kD) is known to bind with high affinity via its N-terminal to β-catenin (85.6 kD). We monitored JF_549_-labeled TCF4 diffusion at a concentration that enabled single PSF localization (40pM) across a titration of unlabeled β-catenin and observed a concentration-dependent decrease in TCF4 diffusion with increasing concentrations of β-catenin (Figure 4C). Interestingly, when a bivalent version of β-catenin was titrated in, TCF4 diffusion was shifted to a minimum of 15.08 µm^2^/s versus 19.39 µm^2^/s when monovalent β-catenin was added, consistent with a slower expected diffusion coefficient of a larger effective particle size. Next we tested whether we could compete away the interaction between TCF4 and β-catenin with the addition of unlabeled TCF4. As expected, diffusion of the JF_549_-labeled TCF4 in complex with β-catenin was increased back to level of JF_549_-labeled TCF4 by the titration of unlabeled competitive TCF4, suggesting competitive disruption (Figure 4D). To demonstrate application of isSMT in the context of drug discovery, a small-molecule inhibitor and a TCF4 competing peptide were tested for their ability to alter the protein motion of JF_549_-labeled TCF4 in complex with β-catenin. Toxoflavin, a known inhibitor of β-catenin, was able to increase the diffusion of JF_549_-labeled TCF4 in a dose-dependent response with a maximum effect similar to the TCF4 competitive peptide (Figure 4D) (Leow et al., 2010). In summary, isSMT represents a new biophysical method for monitoring PPI and protein-ligand interaction with potential sensitivity into the picomolar range.

### OLS is amenable to a variety of SMLM techniques and acquisition schemes

The homogenous illumination, high spatiotemporal resolution, imaging speed and overall robustness of the OLS system is likely to have broad advantages across multiple biological microscopy techniques. To demonstrate the ability to image SMT across 2 spectrally distinct fluorophores, we captured a JF_549_ and a JF_646_-labeled Halo-KEAP1 time series sequentially resulting in clear single-molecule resolution across both wavelengths (Figure 5A). KI-696 was dose titrated and the response was measured across both JF_549_ and JF_646_-labeled Halo-Keap1 to demonstrate the ability to measure changes in protein motion across 2 wavelengths (Figure 5B). Importantly, despite a drop in sCMOS quantum efficiency in the far-red spectrum resulting in lower SNR (Figure S10A), we captured SMT data using the red-shifted JF_646_ and obtained EC_50_ values that were consistent with the brighter JF_549_ dye with measured EC_50_s of 4.96 nM and 6.45 nM for JF_549_ and JF_646_, respectively. Robust measurement of protein dynamics with lower SNR is potentially explained by minimal decrease in lower bound error rate (ERLB) (Figure S10B). Taken together, OLS-based SMT is likely to have applications in multi-color imaging and facilitates imaging of lower quantum yield fluorophores, broadening the palette of available fluorophores to measure protein motion across biological applications.

**Figure 5:**
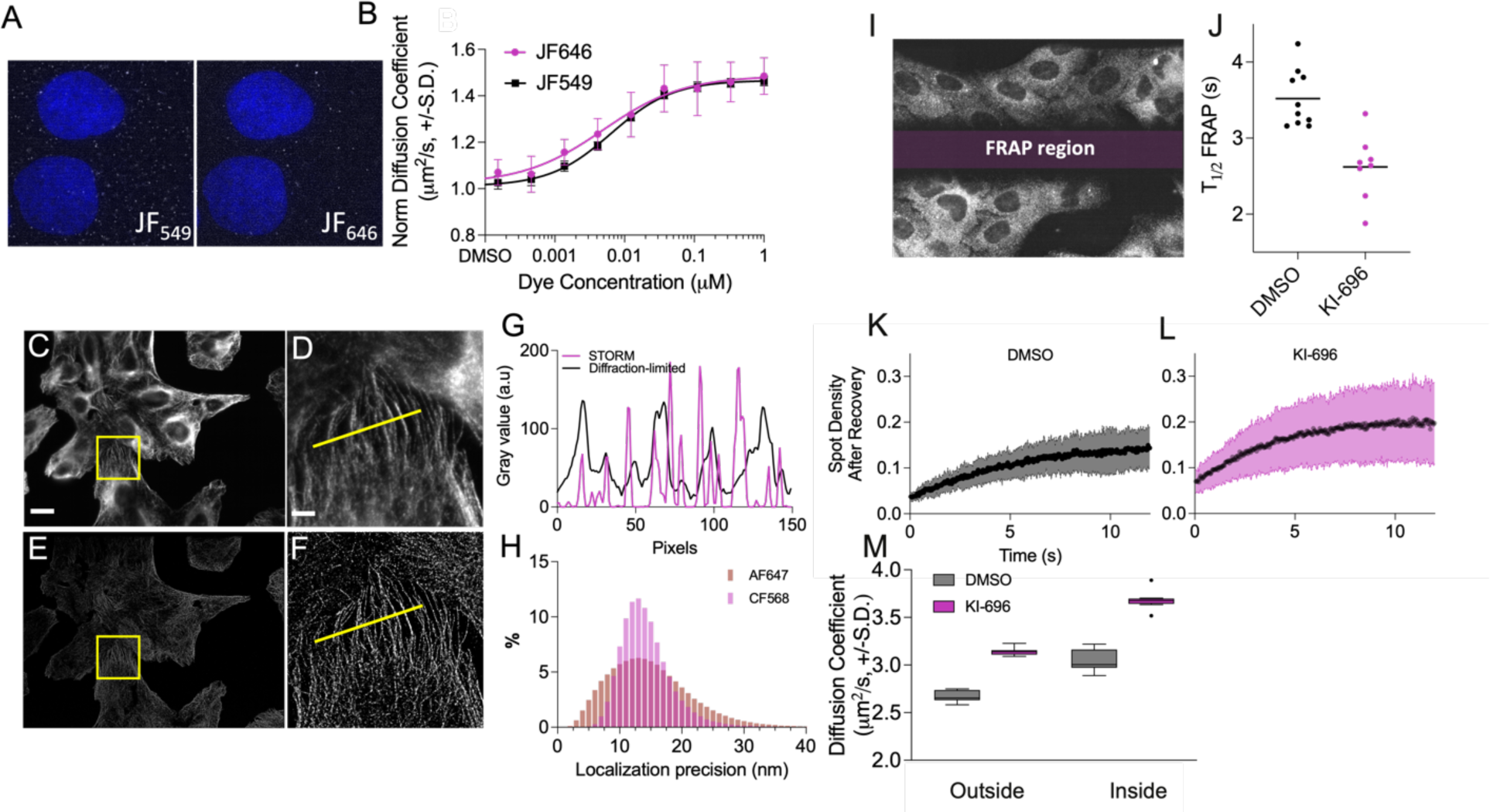
OLS is amenable to a variety of SMLM techniques and acquisition schemes. **(A)** JF_549_ and JF_646_ co-labeled Halo-KEAP1 U2OS cells imaged within the same FOV. **(B)** 10-point dose-response of KI-696-treated Halo-KEAP1 U2OS cells co-labeled with JF_549_ and JF_646_. **(C)** Diffraction-limited image of a full OLS FOV of immunofluorescently labeled tubulin with AF647-conjugated secondary antibody (see methods). Scale bar represents 20 μm. **(D)** Zoomed-in region of interest from (C). **(E)** STORM reconstruction of full OLS FOV, as in (C). **(F)** Zoomed-in region of interest from (E) as in (D). Scale bar represents 5 μm. **(G)** Line profile from yellow lines in (D) and (F) gray value (a.u) to compare spatial resolution of microtubules. **(H)** Localization precision histogram for each of AF647 and CF568 labeled secondary antibodies used to stain microtubules with OLS illumination with a 0.4 ms integration time. **(I)** Representative image of correlative FRAP/SMT where a central region is bleached using the OLS line scan prior to spot recovery after photobleaching. Areas outside and inside of the FRAP region are used to measure SMT. **(J)** T_1/2_ FRAP for Halo-KEAP1 U2OS cells treated with DMSO or 1 μM KI-696 labeled with 400 pM JF_549_-Halo ligand, black line denotes median, and each spot represents an individual FOV. **(K)** and **(L)** Spot density after recovery over time for each of DMSO (K) and 1 μM KI-696 (L). Standard deviation shown in confidence bands for 8-10 FOVs per condition. **(M)** Diffusion coefficient from SMT for 400 pM JF_549_-Halo ligand concentration in bleached (Inside) and unbleached (Outside) region.

Due to the short integration time resulting from the OLS line scan, we investigated whether STORM imaging in fixed cells would result in images with high x,y resolution across a large FOV. Cells labeled with anti-tubulin primary antibodies and AlexaFluor647 (AF647) or CF568-conjugated secondary antibodies were imaged using STORM across a full 60x field of view with a total imaging time of approximately 60 seconds (Figure 5C-F). We observed that, despite the low integration time of 400 μs used in OLS, spontaneous photoswitching provided enough photons to achieve a lateral localization precision of approximately 15 nm using either AF647 or CF568 (Figure 5G and H). Our results demonstrate the potential to conduct high-speed, high-throughput phenotypic screening with STORM or other SMLM techniques with OLS illumination.

Next, we decided to leverage the line scan component of OLS to conduct a correlative SMT-FRAP experiment on KI-696 treated Halo-KEAP1cells labeled with JF_549_. We bleached a 240 × 40 μm region by focusing the scan region on a narrow subset of the FOV for a duration of 100-200 sweeps prior to acquiring the full FOV under normal SMT acquisitions (see Methods). This resulted in each frame containing a bleached (FRAP region) and an unbleached region (Figure 5I). Instead of capturing the intensity recovery after photobleaching over time, the normalized spot density after recovery was measured (Figure 5K and L). We measured an average T_1/2_(DMSO) of 3.52 s (SD= 0.38) and T_1/2_(KI-696) of 2.27 s (SD= 0.43) at a JF_549_ dye concentration of 400 pM that appears optimal to separate the 2 conditions (Figure 5J). These results are consistent with our reported SMT measurements and indicates that KI-696 significantly increases Halo-KEAP1 local protein dynamics. To further ascertain the consistency of our FRAP measurement, we applied SMT to both bleached and unbleached regions of the FOV. In both cases, we were able to measure an increase in Halo-KEAP1 dynamics under a 1 μM KI-696 perturbation at a range of dye concentrations (Figure 5M and S10C). These results highlight and confirm that the advantageous properties brought upon by OLS in combination with the presented tracking algorithm enable sensitive spot detection and SMT even when labeling sparsity is not optimized for.

These data demonstrate that the large FOV, OLS scanning, and high SNR with fast integration times enabled 2 color SMT, STORM and FRAP to be implemented with scale, speed and consistency. We anticipate that our platform will be compatible with other approaches such as fluorescence correlation spectroscopy (FCS) and image correlation spectroscopy (ICS) and could enable leveraging such advanced microscopy methods in high content applications.

## Discussion

We have demonstrated how OLS, as a novel illumination scheme, enhances several properties of SMLM and SMT-based techniques. Compared to previously described illumination methods applied to SMLM and SMT, OLS provides a large and homogenously illuminated FOV, finer sectioning ability, superior SNR, and high spatiotemporal resolution. We demonstrate OLS illumination implemented on 4 distinct microscopes, highlighting robustness and consistency of the systems. The stability and throughput that OLS provides will enable important large-scale studies illustrated by our current ability to screen >300,000 drug-like molecules against drug targets of interest. The improvement in rejecting out-of-focus light enables better single-molecule detection and localization, making OLS suitable for conducting SMT on a variety of cellular systems and protein targets where previously, background fluorescence would be limiting. The data presented in this manuscript allow us to anticipate achieving high SNR SMT results in more complex cellular systems such as spheroid cultures from immortalized cancer cells.

OLS enables data acquisition at frame rates of at least 1250 Hz without compromising SNR and tracking fidelity. As illustrated with Halo-KEAP1, a frame rate of 400 Hz is required to fully characterize the increase in diffusion of drug-induced release from NRF2. We anticipate that there are many biological processes that involve rapid protein motion that were previously unmeasurable with other SMT illumination methods. OLS will provide an opportunity to tackle new protein and cellular mechanisms that are likely occurring at the sub-millisecond scale. Nonetheless, in this current implementation, imaging speed comes at the expense of FOV size, which past a certain size-reduction, will prevent the capture of full cell objects truncating a subset of trajectories. Mitigation strategies using optimized hardware solutions such as cameras with improved speeds are worth further investigating in this area.

Our analysis suggests that the greatest source of variation in SMT measurement stems from cell heterogeneity in protein motion. The large FOV enabled by OLS allows for the simultaneous capture of upwards of 70 U2OS cells in culture, enabling the capture of inter-cell heterogeneity. This was exemplified with PCNA where we were able to simultaneously assign the cell cycle phase and monitor protein dynamics. We show that PCNA protein dynamics are slower during S-phase which correlates to a protein enrichment at sites of DNA replication. During G1, G2 and M phases, we observed a robust increase in dynamics corresponding to the majority of PCNA being homogeneously distributed throughout the nucleus. These findings are consistent with previous characterization of PCNA dynamics (Zessin et al., 2016) but there are important improvements in the approach presented here. Cell cycle was predicted computationally through machine learning models using localization of PCNA as opposed to manual assignment. Additionally, our OLS platform enabled rapid and automated capture and analysis of thousands of cells versus tens of cells per condition typically analyzed using manual SMT approaches. When characterizing protein motion in heterogeneous cell population and potential rare cell sub-types, automated cell classification along with scaled SMT data collection and analysis will be critical to enable work that is comparable with flow cytometry and other single-cell analysis techniques.

Given the versatility of OLS, we were able to apply SMT to directly study protein dynamics in solution. Despite the intrinsic difficulty in generating trajectories for fast-diffusing protein in solution, we validated isSMT measurements by showing a strong overall agreement between experimental measurements and the corresponding Stokes-Einstein theory. We demonstrated the method’s relevance by measuring the interaction between TCF4 and β-catenin. We also demonstrated the ability to leverage isSMT to monitor the disruption of this protein interaction with a competitive peptide and small molecule inhibitors, suggesting that there could be relevant application in drug discovery.

Importantly, isSMT requires picomolar concentrations of purified protein enabling assay development of difficult to purify proteins and the sensitivity to measure sub-nanomolar affinities. We speculate that ligand interactions can alter protein diffusion in ways beyond those caused by disruption of PPIs, including ligand-induced confirmational changes, protein stability changes and perhaps more subtle changes to protein hydration shell resulting in small but measurable changes in diffusion.

We highlighted the versatility of OLS beyond SMT by demonstrating its suitability for rapidly capturing STORM datasets, despite very low illumination and integration times, and achieving a lateral resolution of ∼15 nm for both AF647 and CF568. Finally, we demonstrated a correlative SMT/FRAP approach by which to study protein diffusion. This simultaneous acquisition may enable the measure of protein dynamics at different timescales allowing for the capture of a wider range of protein dynamics comprised of discrete but biologically important subpopulations. As with SMT, we expect OLS to enable large-scale advances in microscopy studies that benefit from high resolution, homogeneous and rapid stroboscopic illumination, and rapid frame rates. We recognize that data storage and processing of large datasets generated by capturing time series of multiple large FOVs can be infrastructurally prohibitive. However, as phenotypic imaging becomes central to systems biology studies and drug screening, we expect an increase in demand for more advanced microscopy technology that enable more sophisticated assays such as OLS.

Here, we provide an initial characterization of the broad utility and ease of implementation of OLS using a series of experimental setups: we benchmark OLS against established methods, demonstrate its utility in SMLM using proof-of-concept experiments, highlight that the OLS platform can provide fundamental insight into biological processes as complex as the cell cycle, expand the technological flexibility of the platform to include FRAP and STORM, and even introduce a novel biophysical method for investigating protein dynamics in solution. However, this minimal set of experiments is not a comprehensive description of the utility of OLS; rather, it is the ease of implementation of OLS, coupled with this experimental flexibility, that will provide the research community with a valuable tool to bring SMLM to bear on many dynamic biological systems.

## Acknowledgements

The authors would like to thank all current and past employees of Eikon Therapeutics, especially Rand Miller and Jesse Vargas for providing critical feedback and editing towards this work as well as Roger M. Perlmutter for helpful discussions and comments.

## Methods

### OLS microscopy

SMT image acquisition for OLS datasets was performed on a custom-built microscope based on a Nikon Ti2, motorized stage, stage top environmental chamber (OKO labs), quadband filter cube (Chroma), custom laser launch with 405 nm, 561 nm, and 642 nm wavelengths, delivering >10 mW, >150 mW, and >150 mW of power to the back focal plane of the objective, respectively. The custom laser launch consists of three externally triggerable free-space laser sources (Cobolt 06-MLD; Huebner Photonics; 2RU-VFL-P-2000-560-M; MBP Communications Inc.; VFL-P-2000-642-M; MBP Communications Inc.).

The Oblique Line Scanning (OLS; see Figure S1A and S1D) unit is attached to the back-port of the microscope providing optical excitation and scanning. The OLS unit receives collimated Gaussian-shaped optical excitation via a polarization-maintaining single-mode fiber coupled to a laser beam coupler. The laser excitation is sent through a combination of a Powell lens, custom-designed cylindrical lenses, and an achromatic lens to shape the beam into a laser line. The beam is guided over a set of two position-adjustable right-angle prisms followed by an aspherized achromatic lens to position the beam and focus the scan-axis onto galvanometric scanning mirrors, which is adjusted to position the beam at an offset of 3.8 mm to the central optical axis in the objective’s back focal plane to achieve an illumination light sheet in the sample of at an inclination angle of 60° (see Figure S1E).

Fluorescence emission was passed through a high-speed filter wheel (Sutter Instruments) and collected with a backlit sCMOS camera (ORCA-Fusion BT, Hamamatsu). The sCMOS camera is operated in progressive mode with an exposure time of 407 µs at an internal line interval of 4.87 µs in order to achieve a virtual rolling slit of ∼200% of the optical excitation and fluorescence line width (see Figure S1F). Images were acquired with a 60x 1.27 NA water immersion objective (Nikon). Environmental chamber was set to 37° Celsius, 95% humidity, and 5% CO_2_.

The default optical excitation is set to an average power of 250 mW at the back focal plane of the objective. Given the 90% transmission of the equipped objective lens for a wavelength of 561 nm, operating the laser at 100 Hz with a duty cycle of 90% for a 100 fps acquisition yields a peak power of 250 mW or a pulse energy of 2.25 mJ/pulse in the sample plane. Assuming a non-scanned line, at an angle of inclination of 60° and with an excitation line adjusted to 280 µm in the focal plane (ie 12.5% wider than the FOV to ensure homogenous and complete coverage), the peak power density and the peak energy density can be estimated at 25.77 kW/cm^2^ and at 231.9 J/cm^2^, respectively. Considering a scanned line at 22.2 µm/ms, the local effective optical excitation per position in the sample can be estimated at 1.85 % of the entire optical excitation by scanning across the entire FOV, which subsequently leads to an instantaneous and local average peak power density and instantaneous and local average peak energy density of 476 W/cm^2^ and 4.29 J/cm^2^, respectively.

System hardware control is realized in a custom-designed and user-configurable electric circuit board for software interfacing, synchronization, and device control. Data acquisition control is realized in a custom-designed and user-configurable acquisition script in MicroManager for raster scanning 384 well plates, a custom-designed automated focusing routine (see Figure S1B). 1 frame of Hoechst and Potomac Red channel were collected at the same frame rate for downstream registration of trajectories to nuclei and cytoplasms respectively.

### HILO Microscopy

SMT Image acquisition for HILO datasets using was performed on a custom-built microscope based on a Nikon Ti2, motorized stage, stage top environmental chamber (OKO labs), quadband filter cube (Chroma), custom laser launch with 405 nm and 561 nm wavelengths, delivering >10 mW and >150 mW of power to the back focal plane of the objective, respectively. Fluorescence emission was passed through a high-speed filter wheel (Finger Lakes Instruments) and collected with a backlit sCMOS camera (ORCA-Fusion BT, Hamamatsu). Images were acquired with a 60X 1.27 NA water immersion objective (Nikon). Environmental chamber was set to 37° Celsius, 95% humidity, and 5% CO_2_. For each field of view, 150 SMT frames were collected at a frame rate of 100 Hz, with a 2 ms stroboscopic laser pulse.

### Signal to noise ratio definition and quantification

Signal to noise ratio (SNR) is defined based on the likelihood ratio for a hypothesis test comparing: a target-absent condition, where the local image is modeled by the sum of a constant offset, and independent Gaussian-distributed noise; and a target-present condition, where the local image is modeled by the sum of a centrally located Gaussian peak (with known width but unknown amplitude), independent Gaussian-distributed noise, and a constant offset. Following the method described in (Serge et al., 2008), the SNR is expressed as

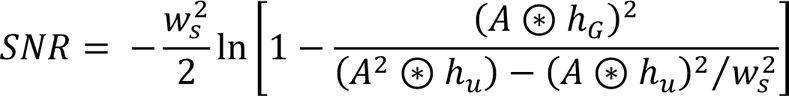

where:

- *A* is the image, cropped to the current region of interest (ROI);
- *w_s_* is the side length, in pixels, of the square ROI;
- ⊛ is the inner product operator;
- *h_G_* is a zero-mean detection kernel, matched to the expected Gaussian target profile, and with ∑*h*_G_^2^ = 1 where the summation is taken over the ROI;
- *h_u_* is a uniform kernel (i.e., it has a value of 1 over the entire ROI).

### Single-molecule tracking methods

Single-molecule tracking (SMT) data were processed with a custom pipeline operating on image sequences produced by the microscope. Briefly, individual emitters were detected by applying a generalized log-likelihood ratio test to every 11×11 subwindow in the image as described above (*Signal to noise ratio definition and quantification*) (Serge et al., 2008). We detected emitters by identifying pixels with a log-likelihood ratio exceeding 14 (for cSMT) or 16 (for isSMT). Detected emitters were localized to subpixel precision in a two-stage procedure. First, the subpixel location was estimated by computing points of maximum radial symmetry (Parthasarathy 2012). Second, this estimate was used to seed an iterative Levenberg-Marquardt fitting routine to a 2D integrated Gaussian within a 11×11 pixel subwindow centered on the detection (Laurence and Chromy 2010; Smith et al., 2010).

Localized emitters were linked in time to produce trajectories using a modification of Sbalzerini’s hill-climbing algorithm that uses Gibbs sampling to estimate data association uncertainty (Hue et al., 2002; Sbalzarini and Koumoutsakos 2005). In all SMT we prohibited links longer than 1.25 µm for cSMT (2.5 µm for isSMT) and links over more than 2 gap frames to limit association error. Emitters were assigned to segmentation categories (nucleus, cytoplasm) by comparing their subpixel location with the semantic masks produced by the segmentation routine.

### Measurements on trajectories

When reporting the number of trajectories, we excluded singlets (trajectories with 1 detection), as these do not contribute information to most dynamical estimates.

Average diffusion coefficients were computed with the mean squared displacement method (*D_est_* = MSD_2D_/4Δ*t*) (Hue et al., 2002; Michalet and Berglund 2012). This estimator is expected to overestimate the diffusion coefficient by *σ_loc_^2^*/Δ*t*, where *σ_loc_^2^* is the variance of the 1D localization error and Δ*t* is the frame interval.

To resolve trajectories in multiple dynamical states, we inferred the coefficients of a Brownian mixture model over a grid of diffusion coefficient values and localization error values using state arrays, a variational Bayesian routine based on the Dirichlet process mixture (Corduneanu and Bishop 2001; David and Michael 2006; Heckert et al., 2022). Mixture components were selected as the Cartesian product of 100 diffusion coefficients log-spaced between 0.01 and 100 µm^2^/s and 31 localization error values from 0.02 to 0.08 µm (1D standard deviations). Occupations are reported as the mean posterior probabilities of each diffusion coefficient marginalized over all values of localization error. To make inference tractable, we limited inference to 10000 trajectories randomly sampled from each well.

Bias estimation in single population samples was performed using analytical calculations that capture the probability of false linking and jump-length-distribution truncation due to a finite search radius. For isSMT, spatial locations are uniformly distributed, allowing the probability of false linking to be estimated based on the observed emitter concentration and a nearest-neighbor linking model. This probabilistic analysis also allows characterization of the false-link jump length distribution. Single-population bias estimates incorporate these false-linking artifacts, and the prohibition on jump-length observations greater than the search radius. As shown in Figure 4A, these bias expressions are used to improve diffusion-coefficient estimation accuracy.

### Empirical estimate of linking precision

To estimate the accuracy of the linking algorithm, we used a bootstrapping procedure. Detections from the first and second halves of the movie were superimposed, and the tracking algorithm was run on the resulting set of detections while blinded to the origin of each detection. From this we computed the fraction of links generated where individual detections were joined from different halves of the movie. Since this fraction neither accounts for erroneous links between detections in the same half of the movie nor for the effects of photobleaching, it forms a lower bound on the linking error rate (ERLB).

### Estimate of SMT dynamic range

To estimate the impact of frame rate on SMT dynamic range (as in Figure 2C and Figure S4), we considered the bounds on the possible values of the mean squared displacement (MSD) estimator of the diffusion coefficient (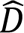) for a Brownian particle with Gaussian localization error. The MSD estimator is biased upwards by localization error according to 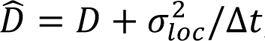, where 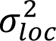 variance of the 1D localization error and Δ*t* is the frame interval. Because *D* is nonnegative, this yields 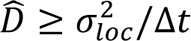. In the opposite limit, when the true jumps of the particle are much larger than the search radius *R* used for tracking, the jumps are uniformly distributed within the tracking range gate (a circle of radius *R*) and the MSD estimator of the diffusion coefficient is capped at *R*^2^/8Δ*t*. Together, this yields the dynamic range estimate 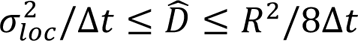. Consequently, the effect of changing frame rate is to translate the dynamic range in the log 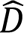 domain. This simple model of dynamic range does not consider the impact of trajectory misconnection (which may change the upper bound) or the non-MSD estimate of diffusion coefficient in state arrays (which may reduce the lower bound).

### Optical-dynamical simulations

To evaluate tracking methodology and the impact of frame rate on SMT dynamic range, we performed optical-dynamical simulations. These simulations use a scalar diffraction approximation to a paraxial imaging system with NA = 1.2 (Hanser 2004). Briefly, we first simulated discrete mixtures of Brownian motions without state transitions at frame rates 12.5, 25, 50, 100, 200, 400, 800, or 1600 Hz in a cube with dimensions 45×45×8 µm^3^ (XYZ). Particles were initialized at density 0.31 or 0.62 particles per cubic micron (depending on the simulation), and were subject to photobleaching at probability 0.03 per frame. Positions of particles coinciding with 500 µs pulses (modeling the stroboscopic illumination in HILO or the rolling shutter in OLS) were accumulated onto a simulated 2D camera via convolution with the system’s 3D point spread function. This yielded a probability distribution of photon arrivals over all simulated camera pixels. Next, we sampled photon arrivals from this distribution as a Poisson process up to mean 90 or 125 photons per particle (depending on the simulation). Finally, Gaussian read noise with RMS 3 photons was added, we multiplied by a 4.3 counts-per-photon gain factor, and the movie was discretized into 16-bit. We then subjected this movie to tracking and state array inference with settings identical to Halo-KEAP1 tracking.

### Assessment of highest source of variance from experiment: Jump resampling experiment

To evaluate the contribution of well-to-well, FOV-to-FOV, and cell-to-cell biases on estimated mean 2D jump length, we acquired a full plate (308 wells) of HILO and OLS data with DMSO-treated KEAP1-HaloTag labeled with JF_549_ at 100 Hz. After excluding the outer ring of wells in the 384-well plate, this yielded a dataset with 308 wells, 12 FOVs per well, and a mean of ∼44 cells per FOV (for OLS) or ∼15 cells per FOV (for HILO). We considered a simple model of jump length. Letting *Y* be the observed 2D jump length, we modeled *Y* as the sum

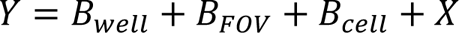

*B*_well_, *B*_FOV_, and *B*_cell_ are random variables modeling bias at the well, FOV, or cell levels, and *X* models the intrinsic stochasticity in jump length conditional on a particular well, FOV, and cell. The simplification is that *B*_well_, *B*_FOV_, *B*_cell_ and *X* are assumed to be independent. Under this simplification, Var(*Y*) = Var(*B*_well_) + Var(*B*_FOV_) + Var(*B*_cell_) + Var(*X*). A more physically realistic model would take into account the potential dependencies between these random variables.

To estimate Var(*B*_well_), Var(*B*_FOV_), Var(*B*_cell_), and Var(*X*), we computed the variance in sample means for four different jump resampling schemes:

1. Sample *N* jumps from the whole plate. The resulting sample mean averages over all sources of variability (*B*_well_, *B*_FOV_, *B*_cell_ and *X*).
2. Sample a well, then sample *N* jumps from that well. The resulting sample mean averages over *B*_FOV_, *B*_cell_, and *X*, and the variance over these sample means is expected to approach Var(*B*_well_) as *N* becomes large.
3. Sample a well, sample an FOV from that well, then sample *N* jumps from that FOV. The resulting sample mean averages over *B*_cell_ and *X*, and the variance over these sample means is expected to approach Var(*B*_well_) + Var(*B*_FOV_) as *N* becomes large.
4. Sample a well, sample an FOV from that well, sample a cell from that FOV, then sample N jumps from that cell. The resulting sample mean averages over *X* only. The variance over these sample means is expected to approach Var(*B*_well_) + Var(*B*_FOV_) + Var(*B*_cell_) as *N* becomes large.

1000 rounds of sampling were performed for each sampling scheme and the variances of the biases *B*_well_, *B*_FOV_, and *B*_cell_ were estimated by the difference between the variances over sample means produced by each resampling scheme. As expected, only Var(*X*) depended on the sample size *N* and the other sources of variability were stable with respect to the sample size after ∼100 jumps (Figure S7).

### His-Halo

His-Halo was diluted to 50 pM in imaging buffer consisting of 25 mM HEPES pH 7.4, 150 mM NaCl, 1 mM DTT, 0.03% (w/v) BSA, 0.01% NP-40, with glycerol levels spanning 0 to 40%. Dilutions were transferred to a 384-well plate, which was incubated at 37°C for 15 minutes prior to imaging using OLS. We captured 22-44 well replicates per concentration and acquired 4 FOVs per well at 200 frames per second, and the glycerol titration experiment was performed on two plates using the same protein preparation viscosity calculations are performed following previously described methods (Cheng 2008; Volk and Kähler 2018).

### B-catenin/TCF4

GST-B-catenin (dimer B-catenin) was made via bacterial expression. GST tag was removed via thrombin cleavage and purified by SEC to obtain monomer B-catenin.

His-Halo-TCF4 (1-53) was made via bacterial expression, then labeled with JF_549_ and purified by SEC. Final concentration was determined via Nanodrop and gel band intensity imaged on Cy3 channel of ImageQuant800.

Halo-TCF4 and B-catenin were diluted in imaging buffer (25 mM HEPES-K pH=7.6, 0.1 M EDTA pH=8.0, 12.5 mM magnesium chloride, 100 mM potassium chloride, 0.2 mM PMSF, 1 mM DTT, 0.3 mg/ml BSA, 0.01% NP40, and 30% glycerol, pH=7.9) to final concentrations of 40 pM and 30 nM, respectively. Compounds were printed onto a 384 Cell Vis glass-bottom plate via Echo 655, then incubated with B-catenin for 10 minutes at room temperature prior to addition of Halo-TCF4. Protein solutions were dispensed using an Integra VIAFLO. Plates were then incubated at 37°C for 2 hours prior to imaging.

### STORM and PALM datasets

U2OS cells were fixed in 4% PFA for 10 minutes, washed 3 times with PBS, permeabilized with 0.1% Triton-X for 10 minutes and washed again 3 times with PBS. Cells were then blocked in 2% BSA for 1 hour at room temperature followed by overnight incubation with anti-alpha tubulin antibody (ab7291) at 4 °C. The primary antibody was removed by washing 3 times with PBS and cells were blocked again with 2% BSA for 1 hour at room temperature. Alexa Fluorophore 647-conjugated secondary antibodies were diluted 1:1000 in blocking buffer and incubated for 2 hours at room temperature. Cells were then washed 3 times with PBS to remove excess secondary antibody. Hoechst 33342 was added at 7 uM concentration with the secondary antibodies for nuclei visualization.

Photoswitching buffer for STORM imaging was prepared as previously described (Dempsey et al., 2011) Buffer A was prepared with 0.5 mL 1M Tris (pH 8.0), 0.146 g NaCl and 50 mL of H_2_O. Buffer B with 2.5 mL 1 M Tris (pH 8.0), 0.029 g NaCl, 5 g Glucose and 47.5 mL of H_2_O. GLOX solution was prepared by mixing 14 mg Glucose Oxidase (G2133) and 17 mg/mL of Catalase (C40) with 200 µL of Buffer A. The final photoswitching buffer was prepared by combining 100 μL 1M MEA (M9768) with 10 μL of GLOX and 1 mL of buffer B. This buffer was added to cells in the 384 well plate prior to imaging.

Low laser power (300 mW) was used to capture diffraction-limited images and identify the pertinent focal plane. Laser power was increased to achieve ∼500 mW at the back focal plane in order to initiate photoswitching. 500-5000 frames were acquired, with an integration time of 400 μs.

### Line FRAP acquisition

FRAP datasets U2OS-KEAP1 were recorded by consecutively imaging 5 pre-bleaching frames, bleaching a subregion of the imaging FOV, and capturing fluorescence recovery at an operating framerate of 25 fps. Bleaching a FOV subregion was achieved by scanning the excitation laser for 100-200 sweeps across a subregion of the FOV (16-25% FOV height, 100% FOV width) at high laser power (300 mW) in order to bleach the local fluorescence. Fluorescence recovery was acquired with 400-500 frames following the bleaching step at low laser power (70 mW). Modulating the laser power was achieved by implementing an Acousto-Optical Tunable Filter in the laser engine module (AOTF-Ed 2018-1; Opto-Electronic) adjusted for a bleaching and imaging power of the system’s 560 nm-excitation.

### Cell line engineering of CRISPR Knock-in Halo-tagged proteins

(All U2OS cells were cultured in Dulbecco’s modified Eagle’s medium (DMEM) (Gibco) supplemented with 10% fetal bovine serum (Corning), 100 units/mL penicillin, 100 μg/mL streptomycin (Gibco), at 37 °C and 5% CO_2_.)

To generate KEAP1-Halotag and Halotag-PCNA cell lines, ribonucleoprotein (RNP) complexes included sgRNAs targeting either N- or C-terminal region (Integrated DNA Technologies - IDT) and Cas9 protein (PNA bio, Cat #CP01) were transfected together with linear dsDNA donors (IDT) using Lonza nucleofection method. Each donor consists of 200-300 bp homology arms specific for each target, codon optimized Halotag sequence, and TEV linker (ENLYFQG) between the target and Halotag.

After transfection, the cells were incubated with Halo ligand JF_646_ (Internal) and imaged with ImageXpress system (Molecular Device) to confirm HaloTag integration. Cells were then subjected to single-cell sorting into 384-well plates. Clonal cells were expanded imaged with the ImageXpress system and genotyped by Sanger sequencing to confirm homogenous HaloTag integration.

### Western blot analysis

Protein was extracted from 1-2 million cells by lysing with 1x Cell Lysis Buffer (CST, #9803) diluted in UltraPure Sterile Water (Intermountain Life Sciences, 20804225) with 1x Halt Protease and Phosphatase Inhibitor Cocktail (Thermo Fisher Scientific, 78440). Samples were collected using cell scrapers (FisherScientific, 08-100-241) and placed on ice for 15 minutes and then centrifuged at 15,000 RPM for 10 minutes. The supernatant was transferred to a new tube. Protein concentration was determined with the Pierce BCA Protein Assay Kit (Thermo Fisher Scientific, A55864) according to the manufacturer’s protocol. Samples were run on the automated western blot system Jess (ProteinSimple, 004-650) according to the manufacturer’s protocols. The 12-230 kDa separation module was used for PCNA (CST, D3H8P, Rabbit, 1:100 dilution, #13110), b-Actin (CST, D6A8, Rabbit, 1:50 dilution, #8457S) and Halo (Promega, Mouse, 1:10 dilution, #G9211) detection. Samples were diluted to 0.3 mg/mL with 0.1x Lysis Buffer and 5x Master Mix and heat denatured at 95 °C for 5 minutes and all primary antibodies were diluted with Antibody Diluent 2. All reagents were loaded to a microplate and a 13 capillary cartridge was loaded onto the Jess western blot instrument. Run parameters were set using the COMPASS software v6.0.0 (ProteinSimple). Each protein’s corresponding band was visualized using the COMPASS software and bands were normalized to the loading control b-Actin.

### Cell proliferation

6-well plate cell proliferation – Cells were seeded into 6-well plates (Fisher Scientific, 07-200-83) 150,000 cells/well and left at ambient temperature for 20 minutes to ensure homogeneous settling. The plate was imaged with the Incucyte (Sartorius) taking 9 images per well every 4 hours and phase masking algorithms were applied to determine cell confluence using the Incucyte software v.S3 2019A.

### Sample preparation in 384-well plate

Cells were seeded in tissue culture treated 384-well glass-bottom plates (Cellvis #1.5 cover glass) at 4000-6000 cells per well and allowed to adhere and incubate overnight at 37 °C and 5% CO_2_. Cells were labeled for SMT using JF_549_ and JF_646_ along with Hoechst 33342 and organelle-specific dyes for an hour prior to washing with DPBS (3x) and Fluorobrite DMEM media (2x).

### Cell Segmentation

Segmentation models were trained to segment specific compartments from the Hoechst and Potomac Red images. The models were trained using a U-Net architecture with the following class-balanced entropy loss (Ronneberger et al., 2015; Cui et al., 2019).

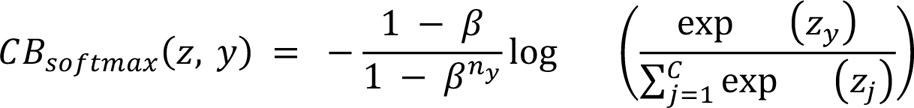

- Z is the predicted output of the model for all classes
- C is the total number of classes
- b is a hyperparameter between 0 and 1
- n_y_ is the number of training samples per class y

For the Hoechst model 3 classes were used representing nuclei, nuclear border, and autofluorescent artifacts. For the Potomac red model 5 classes were used representing nuclei, cytoplasm, nuclear border, cell border, and autofluorescent artifacts.

### Cell cycle classification model

A segmentation model was trained to identify nuclear regions. The input is an image of JF_549_-labeled PCNA and the model generates pixel-level classifications. In addition to identifying and generating mask objects for the nuclei, each nucleus is categorized as being within one specific phase of the cell cycle. From fluorescently labeled PCNA images we were able to categorize the nuclei using the following labels: Mitotic (M), Growth 1 (G1), Early S, Mid S, Late S (S), or Growth 2 (G2).

To evaluate the cell cycle prediction model, we compared the nuclei categorical predictions to manually labeled data (Figure S8). Once the pixels that correspond to each category have been identified we can compute tracking metrics within each mask object.

## Supplementary Figures

**Figure S1:**
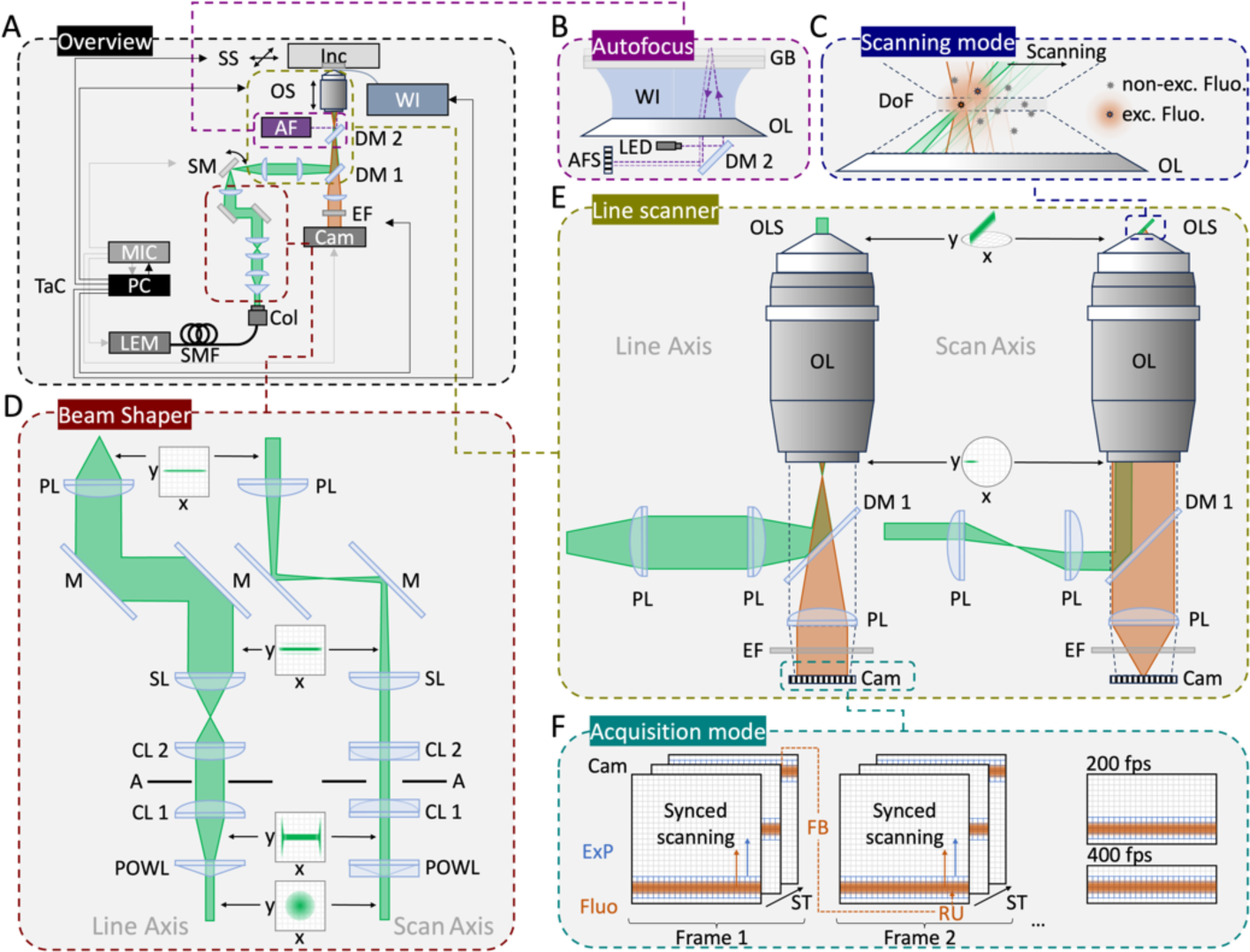
Schematic of Oblique Line Scanning microscope for single-molecule tracking. **(A)** Schematic of the OLS microscope based scanning an inclined excitation light sheet using galvanometric scanning mirrors across a sample placed in an inverted microscope. The OLS microscope is based on a multiwavelength optical excitation provided by a Laser Engine Module and coupled to the beam shaper by a collimator-coupled single-mode fiber. The beam shaper transforms the incoming gaussian-shaped optical excitation to an optical light-sheet that is focused into the back focal plane of the microscope’s objective lens along the light sheet’s line axis and scanned along the scan axis using galvanometric mirrors. The resulting oblique light sheet is sent into a water-immersion-coupled and environmentally-controlled sample holding plate, whereas the relative position of the microscope’s focal plane is controlled by an autofocus unit. The excited fluorescence is spectrally filtered from the excitation light by dichroic filters and emission filters and projected onto a high-speed sCMOS camera. Synchronization of optical excitation, scanning, and acquisition is achieved by a custom-build control unit. **(B)** The autofocus unit is based on detecting a 780 nm-LED reflection on the top surface of the sample-holding glass bottom and repositioning the objective lens to ensure appropriate focal plane positioning within the sample. **(C)** The optical confocal scanning mode is achieved by scanning an inclined and focused light sheet through the objective’s focal plane. Background suppression is achieved by confocal arrangement of the inclined light sheet (green), the objective’s depth of field, and synchronized rolling shutter detection (orange). **(D)** The beam shaping subassembly projected along the line axis (x) and the scan axis (y) shapes collimated optical excitation into a light sheet by a series consisting of a Powell lens, cylindrical lenses, a spherical lens, and a planoconvex lens before encountering the **(E)** The optical line scanning framework is based on an inclined light sheet in the sample plane achieved by focusing the optical excitation along the line axis in the objective’s back focal plane and positioning the optical excitation along the scan axis at an offset position relative to the optical axis of the objective. The corresponding detection of optically-aligned fluorescence is projected onto the camera sensor. **(F)** The OLS acquisition mode relies on detecting fluorescence by matching the camera’s area of exposed pixels and synchronizing the camera’s rolling shutter to the optically-projected intensity line of fluorescence excited by the inclined light sheet. intensity line of fluorescence excited by the inclined light sheet. Abbreviations: A, Aperture; AF, Autofocus; AFS, Autofocus sensor; Cam, Camera; CL, Cylindrical lens; Col, Colimator; DM, Dichroic mirror; DoF, Depth of field; EF, Emission filter; ExP, Exposed Pixels; FB, Fly-Back; Fluo, Fluorescence; GB, Glassbottom; Inc, Incubator; LED, Light emitting diode; LEM, Laser engine module; M, Mirror; MIC, Microscopy illumination control; OL, Objective lens; OLS, Oblique line scan; PC, Personal computer; PL, Planoconvex lens; POWL, Powell lens; RU, Run-up; SL, Spherical lens; SM, Scanning mirror; SMF, Single-mode fiber; SS, Sample stage; ST, Scantime; TaC, Trigger and control; WI, Water immersion

**Figure S2:**
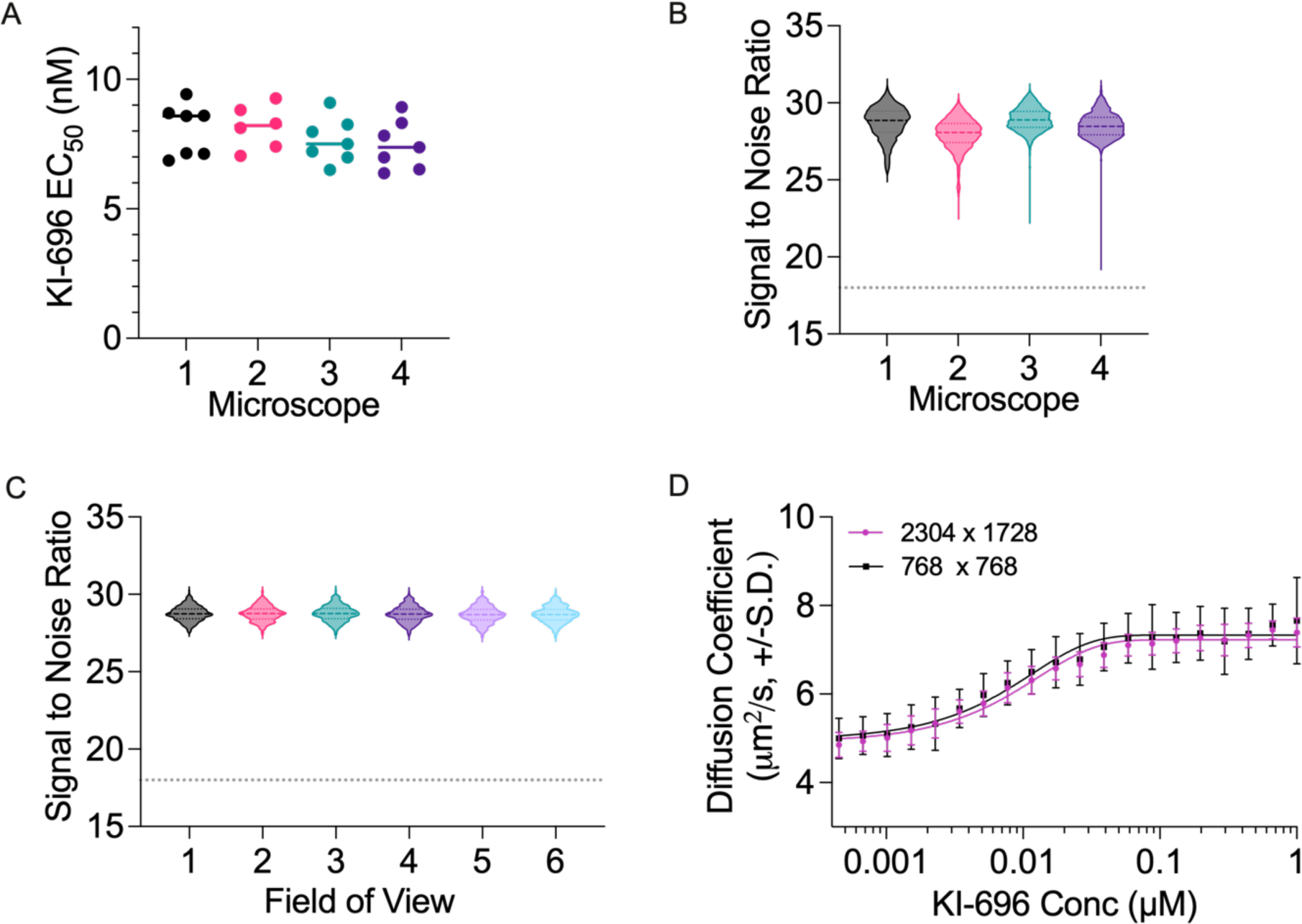
OLS enables reproducible and robust SMT measurements. **(A)** EC_50_ values calculated from each averaged dose-response curve per plate per microscope, black lines represent the median EC_50_. **(B)** Violin plot of the signal to noise ratio (SNR) per microscope, heavy dashed line represents median value. **(C)** Violin plot of the SNR as a function of FOV position within an acquired well. **(D)** 20-point dose-response curve for Halo-KEAP1 U2OS sampled at full OLS FOV (purple) versus a cropped FOV of 768 × 768 pixels (black) representative of a HILO-sized FOV. Error bars denote standard deviation between FOVs.

**Figure S3:**
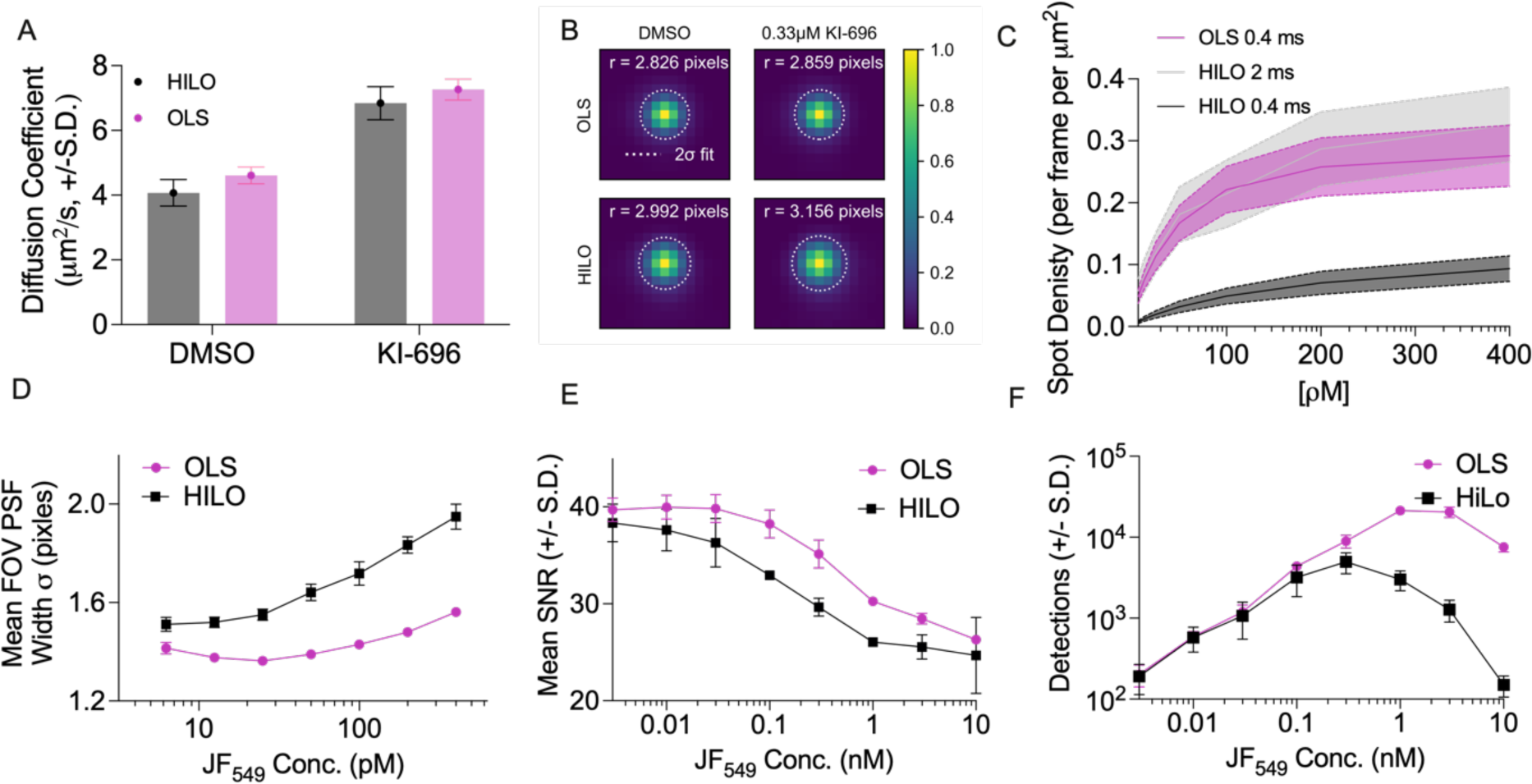
Comparison of motion-induced blurring and confocality between OLS and HILO illumination. **(A)** Measurement of diffusion coefficients for Halo-KEAP1 treated with DMSO or 1 μM KI-696 across 72 FOVs from 12 individual wells for HILO and OLS. **(B)** Estimated point spread functions, calculated by averaging all detections in a representative 150-frame acquisition. The following number of PSFs for each condition: n= 123,596 (OLS-DMSO), n= 3,897 (HILO-DMSO), n= 113,276 (OLS 0.33μM KI-696), n= 13,620 (HILO 0.33 μM KI-696) from one representative FOV. **(C)** PSF detections as a function of integration time. **(D)** PSF width measurement as a function of JF_549_ measured in Halo-KEAP1 cells treated with 1 μM KI-696. **(E)** Mean SNR plotted as a function of JF_549_ measured in Halo-KEAP1 cells treated with 1 μM KI-696. **(F)** Number of spot detections plotted as a function of JF_549_ measured in increasing concentrations of Halo-JF_549_ in solution.

**Figure S4:**
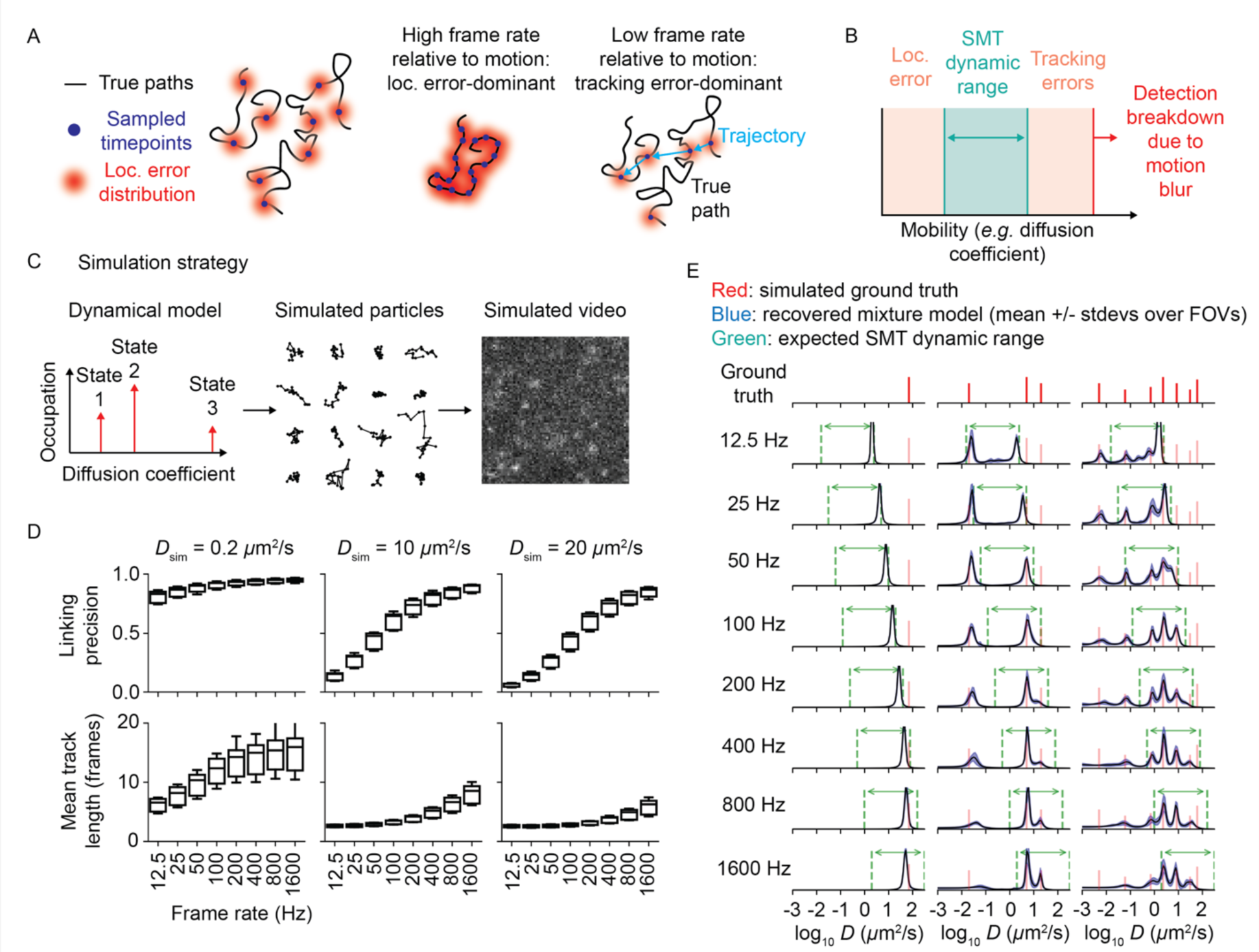
Frame rate determines SMT dynamic range. **(A)** Schematic describing the role of localization and tracking errors for a hypothetical fast moving protein. The rolling shutter in OLS captures the position of dye molecules at discrete timepoints. If these timepoints are too close, the apparent motion is dominated by localization error. If the timepoints are too far apart, reconstructing trajectories becomes challenging and is dominated by misconnections. **(B)** Schematic describing the dynamic range of SMT, bounded on one end by localization error and on the other by tracking errors. An approximation of this range for Brownian motion is 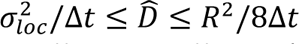 where *σ*^2^ is the localization error variance, Δ*t* is the frame interval, *R* is the search radius, and *D* is the diffusion coefficient (Methods). **(C)** Schematic of simulation approach to test the role of frame rate. Movies were simulated with real-world effects including defocus, motion blur, shot noise, and read noise. (D) Effect of frame rate on linking precision and track length. Linking precision is defined as the fraction of links made by the tracking algo that are correct; track length is the number of points in each trajectory. Quantiles are over simulated movies. **(E)** State array posterior mean occupations for three simulated dynamical mixtures at increasing frame rate. Red lines correspond to the simulated discrete mixture model, blue lines to the state array posterior means, and green lines to the expected SMT dynamic range as defined in (B). Ten simulation replicates were included for each condition.

**Figure S5:**
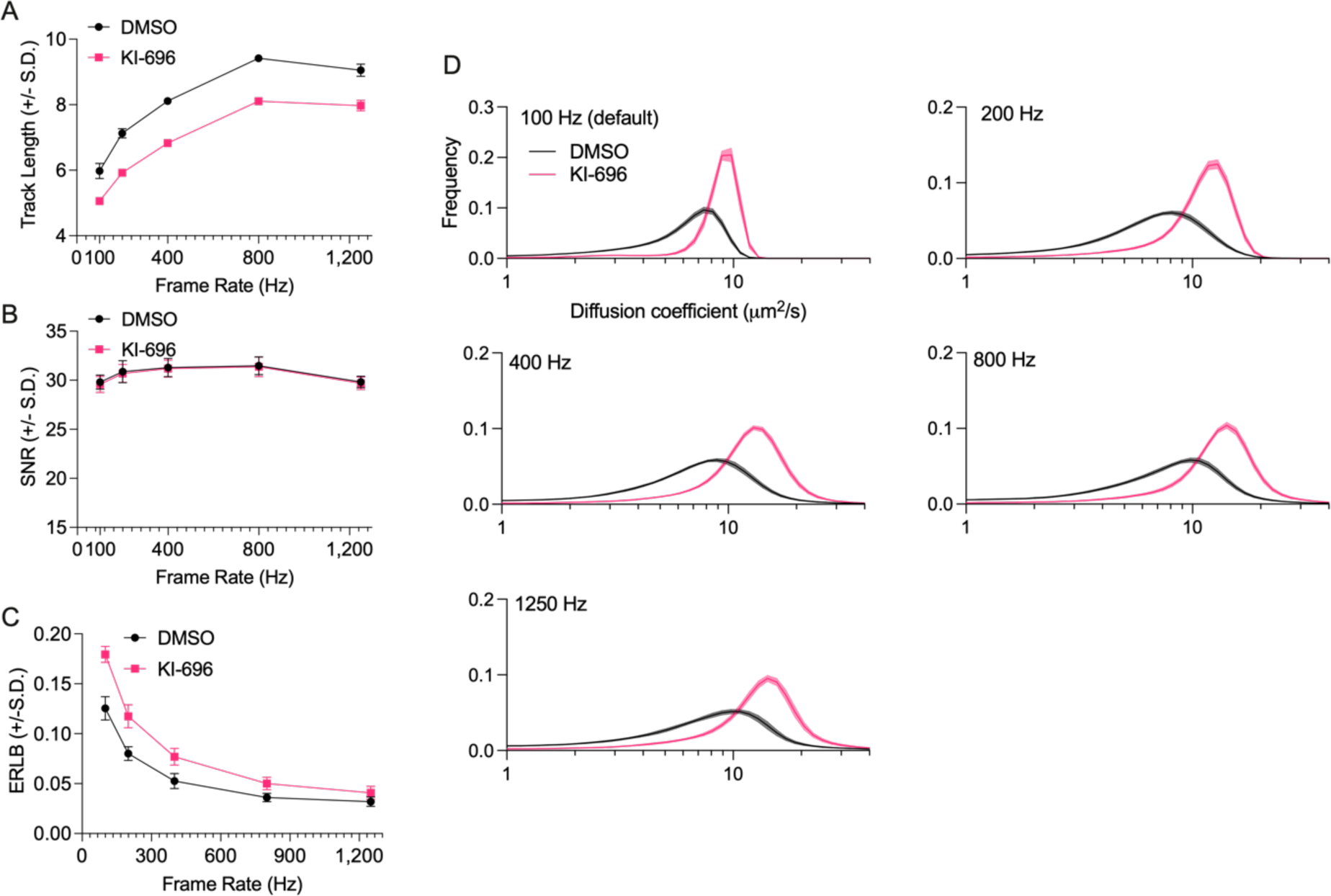
Tracking characterization for KEAP1-HaloTag JF_549_ SMT in U2OS cells with varying frame rates. **(A)** Mean trajectory length plotted as a function of frame rate. Track length is defined as the number of spots linked per trajectory. **(B)** Mean SNR plotted as a function of frame rate, for SNR calculation see Methods. **(C)** mean ERLB plotted as a function of frame rate, ERLB is defined as in “Empirical estimate of linking precision” (Methods). **(D)** State array analysis plotted as a function of frame rate comparing DMSO and 1 μM KI-696 treated Keap1-HaloTag U2OS cells. The number of FOV replicates per frame rate was as follows: n=88 (100 Hz), n=88 (200 Hz), n=132 (400 Hz), n=198 (800 Hz), and n=264 (1250 Hz). Lines are the mean values over all FOVs in the corresponding condition and error bands are the FOV-level standard deviations.

**Figure S6:**
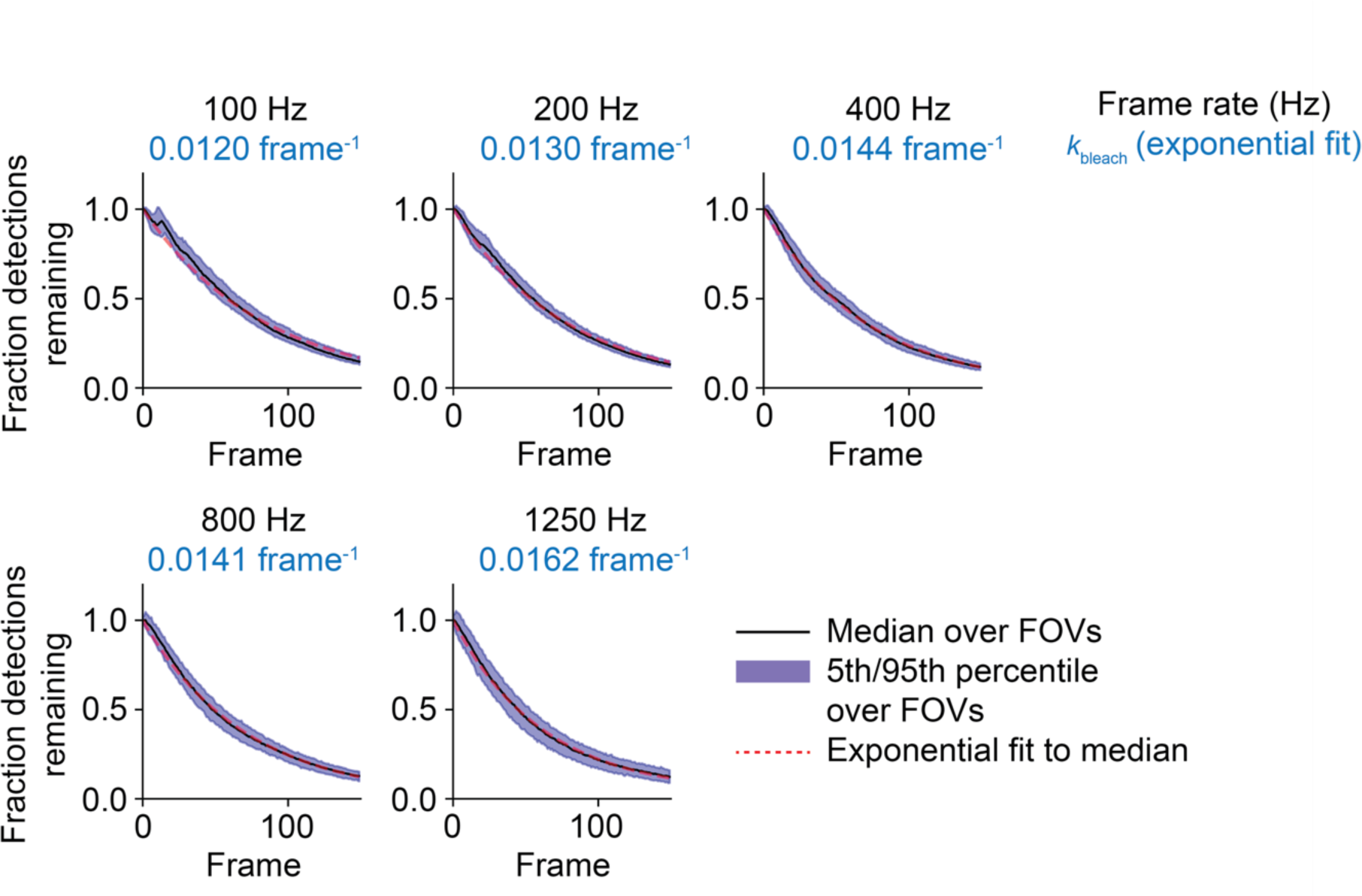
Evaluation of bleaching rate in KEAP1-HaloTag SMT at variable frame rates. Fraction of detections remaining was plotted across frame rate for a given time series. The fraction of detections remaining was defined as the number of detections in each frame divided by the number of detections in the first frame. Exponential fits (blue text below frame rate) were performed with respect to the model *f*(*t*) = *c*_0_ + (1 − *c*_0_)*e*^−*kt*^, where *t* is frame index, *k* is bleaching rate, and *c*_0_ is the unbleached fraction, using an iterative least-squares routine. The number of FOV replicates per frame rate was as follows: n=88 (100 Hz), n=88 (200 Hz), n=132 (400 Hz), n=198 (800 Hz), and n=264 (1250 Hz).

**Figure S7:**
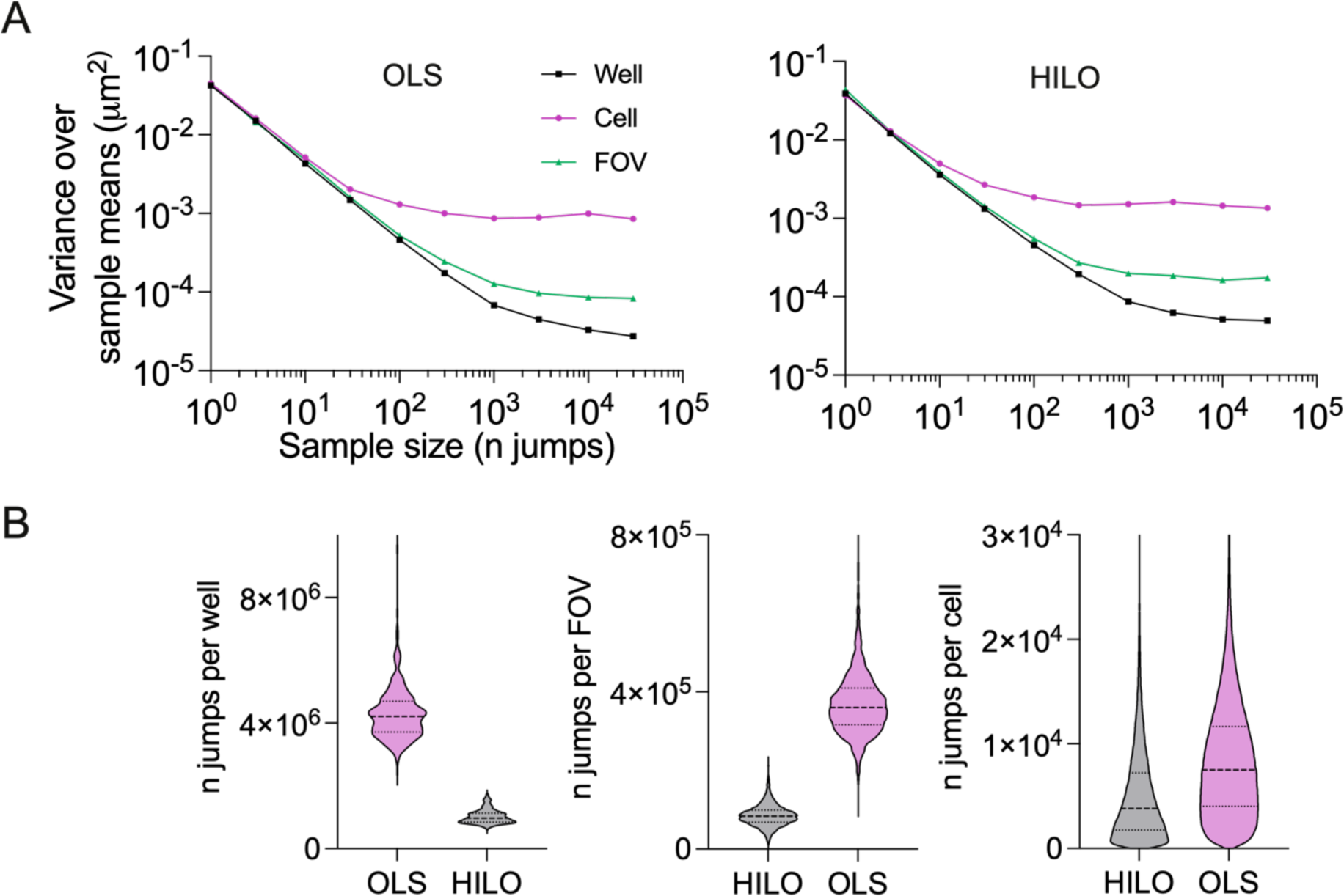
Contribution of well-to-well, FOV-to-FOV, and cell-to-cell biases to 2D jump length, assessed using jump resampling. **(A)** Variance over sample means as a function of sample size for different resampling procedures. A straight line with slope -1 is the expectation from the law of large numbers; sublinearities are due to residual variances over wells, FOVs, or cells. **(B)** Number of jumps per well, FOV, or cell used in these analyses.

**Figure S8:**
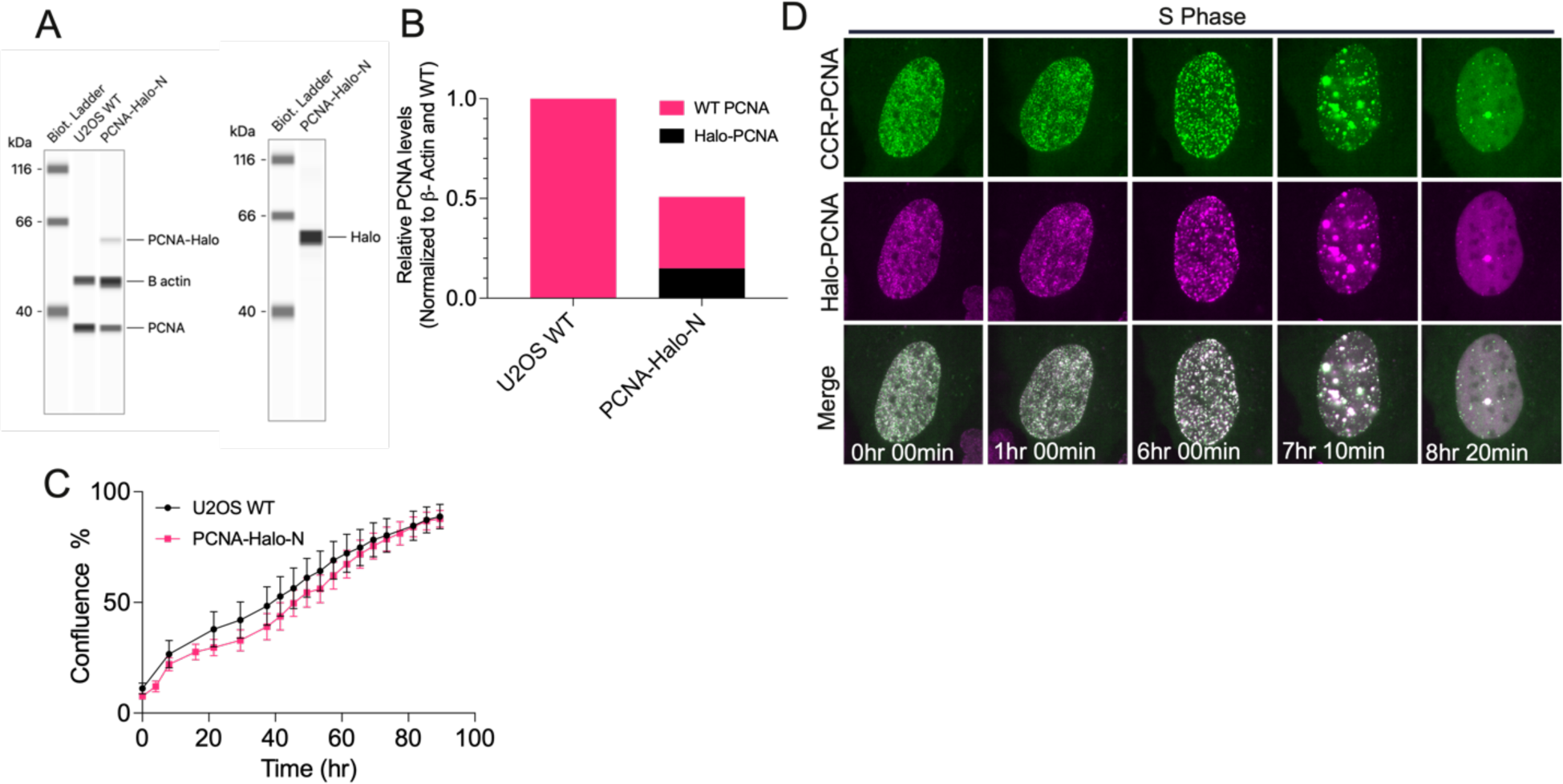
PCNA cell line validation using Western blot and cell proliferation analysis. **(A)** Capillary-based Western blot comparing WT U2OS and N-terminal tagged heterozygous PCNA clone with anti-PCNA antibody (left) and anti-Halo antibody (right). **(B)** Relative WT and Halo-tagged PCNA levels normalized to β-actin in WT and Halo-edited U2OS cells **(C)** Growth curve of WT U2OS and N-terminal Halo-Tagged PCNA. **(D)** Cells labeled with both JF_549_ and CCR PCNA to measure spatial colocalization between the two labels over the course of the cell cycle.

**Figure S9:**
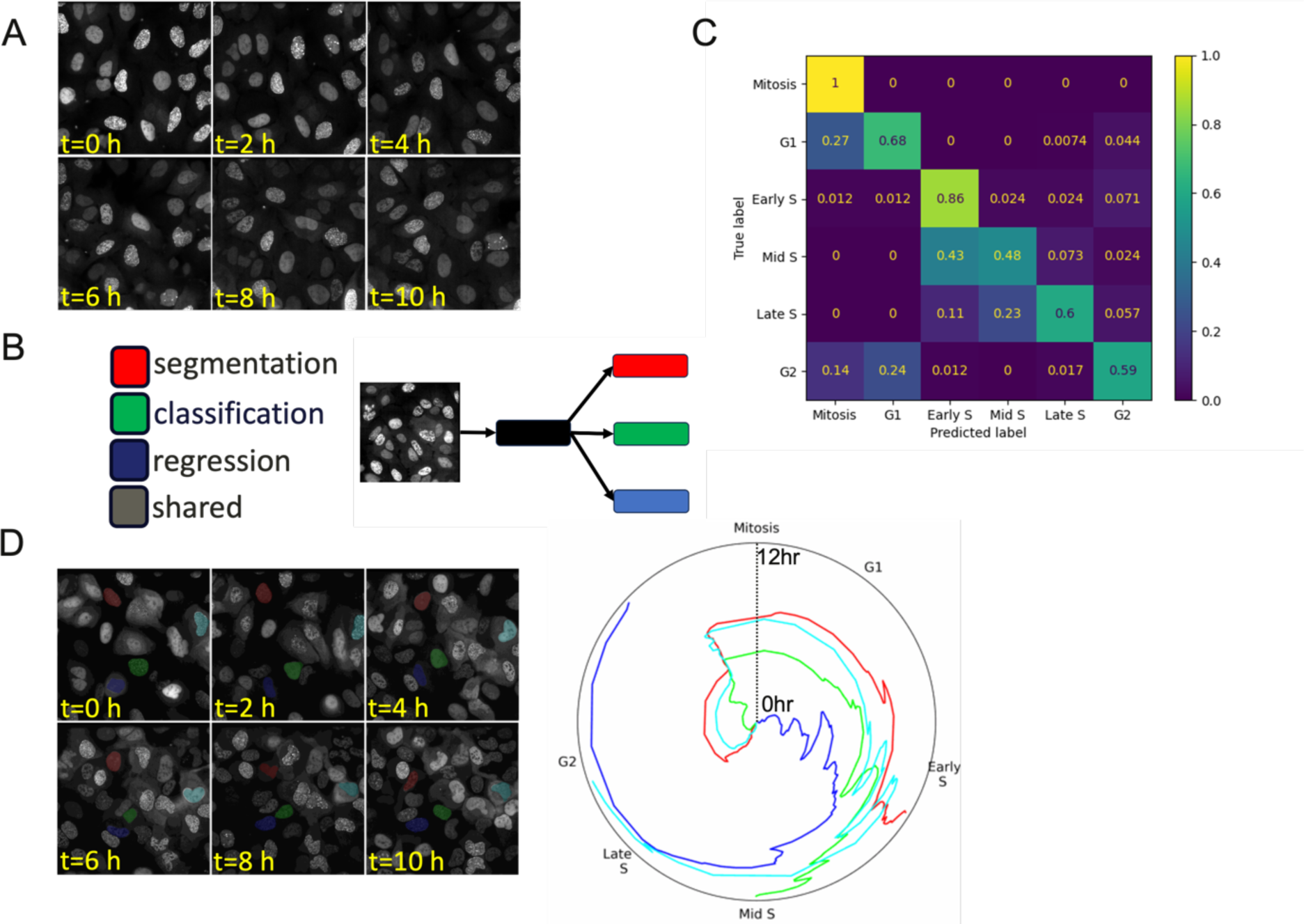
Characterization of PCNA-based cell cycle prediction model. **(A)** Example images from time-lapse of Halo-PCNA captured on OLS with 5 min frame interval for 12 hours. **(B)** Schematic of neural network trained to simultaneously perform segmentation of nuclei, cell cycle classification of nuclei, and cell cycle regression of nuclei. **(C)** Confusion Matrix describing the performance of PCNA-based cell cycle classification. **(D)** Representative time-lapse image of cell cycle progression over a 12 hour window for 4 selected cells (left) with a graph of regression-based prediction of cell cycle progression plotted with a 5 frame moving average (right).

**Figure S10:**
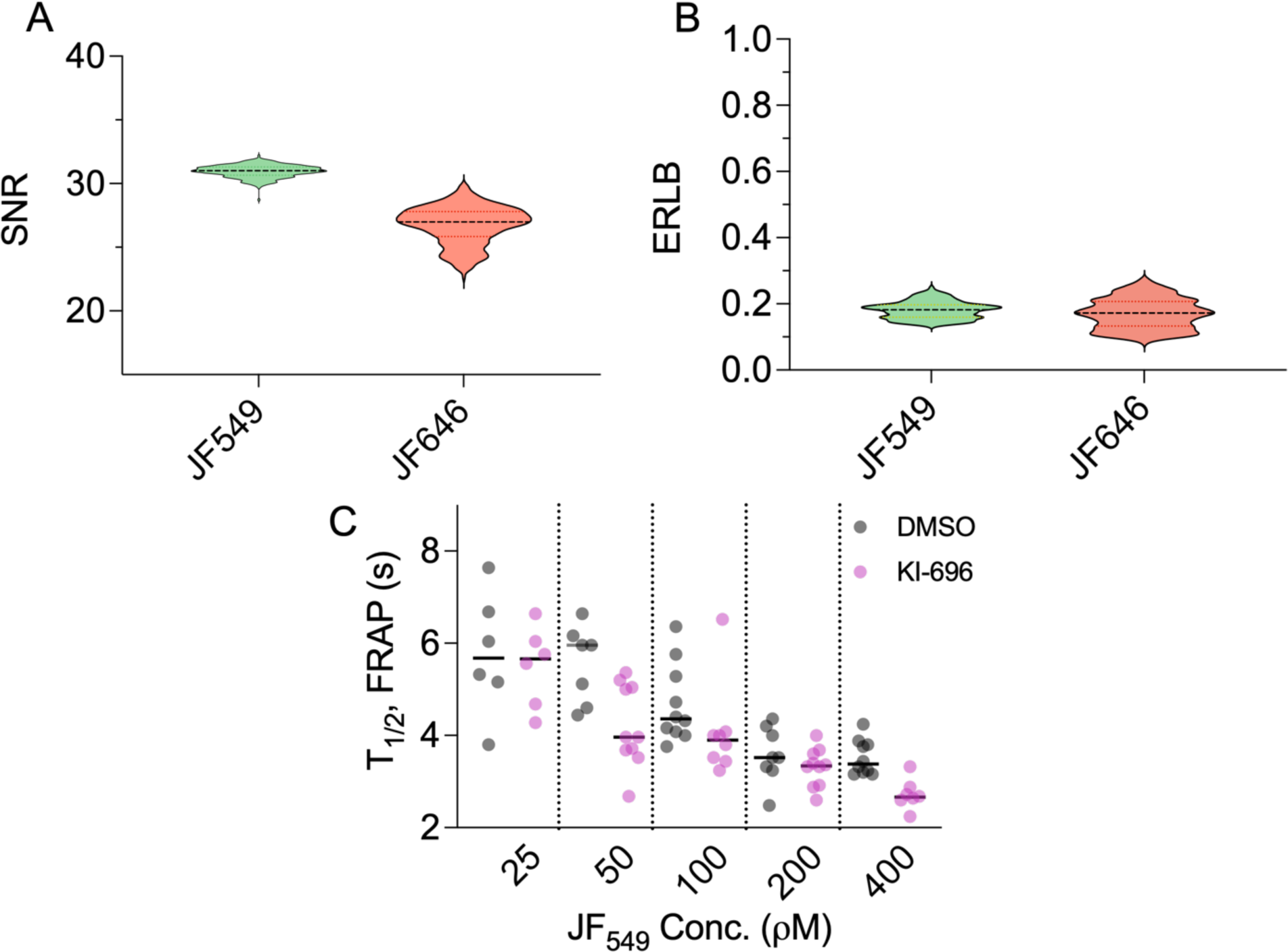
Characterization of dye performance and FRAP with increasing dye concentration. **(A)** SNR comparison between JF_646_ and JF_549_. **(B)** ERLB comparison between JF_646_ and JF_549_. **(C)** Sampling T_1/2_ FRAP measured in bleached region as a function of dye concentration for 6-10 FOVs for DMSO and 1 μM KI-696.

